# The establishment and transmission of novel foraging techniques indicates a capacity for culture in bumblebees (*Bombus terrestris*)

**DOI:** 10.1101/2022.10.10.511574

**Authors:** Alice D. Bridges, HaDi MaBouDi, Olga Procenko, Charlotte Lockwood, Yaseen Mohammed, Amelia Kowalewska, José Eric Romero González, Joseph L. Woodgate, Lars Chittka

## Abstract

The astonishing behavioural repertoires of social insects have been thought largely innate, but these insects have repeatedly demonstrated remarkable capacities for both individual and social learning. Using the bumblebee *Bombus terrestris* as a model, we developed a two-option puzzle box task and used open diffusion paradigms to observe the transmission of novel, non-natural foraging behaviours through populations. Box-opening behaviour spread through colonies seeded with a demonstrator trained to perform one of the two possible behavioural variants, and the observers acquired the demonstrated variant. This preference persisted among observers even when the alternative technique was discovered. In control diffusion experiments that lacked a demonstrator, some bees spontaneously opened the puzzle boxes but were significantly less proficient than those that learned in the presence of a demonstrator. This suggested that social learning was crucial to proper acquisition of box-opening. Additional open diffusion experiments where two behavioural variants were initially present in similar proportions ended with a single variant becoming dominant, due to stochastic processes. This could lead to the emergence and maintenance of local cultural variation. These results suggest that bumblebees, like mammals and birds, may have the capacity for culture and to sustain cultural variation.

## Introduction

The diversity of behaviours observed in some insect societies is on a par with, or exceeds that, of some mammals (1,2), and includes the construction of architecturally complex, climate-controlled nests, and the division of labour between foraging, brood care and nest defence (3,4). Indeed, outside the human realm, their nesting structures are unparalleled in terms of their regularity, sophistication, and their scale in proportion to body size (5). There is profound variation in foraging specialisations, architectures and social organisations not just between related species of social insects, but more intriguingly, even within species (4,5). However, unlike in humans, these specialisations have historically been viewed as a limited set of pre-programmed responses to external stimuli resulting from evolutionary trial-and-error processes (2). However, the innate repertoire of these insects is supplemented by a remarkable capacity for learning that has been recognised for decades. The acquisition of the honeybee dance language is, perhaps, the best-characterised example of social learning described thus far in an invertebrate (6), and as early as 1884, Charles Darwin suggested that “nectar-robbing” of flowers by bumblebees could spread socially. Here, a forager bites into the base of flowers to extract the nectar, which does not pollinate the plant (7). That socially-transmitted nature of this behaviour has since been confirmed: in the wild, nectar-robbing is thought to have repeatedly arisen as independent innovation events, and to spread through local bumblebee populations via rapid social learning (8,9).

The spread of nectar-robbing in the wild is of particular note because it bears strong similarities to another “larcenous” behaviour: milk bottle-opening by tits (*Parus* spp., 10,11). Here, tits across the UK learned to take advantage of a novel, energy-rich food source in their environment by pecking the lids of milk bottles left on people’s doorsteps to access the cream at the top (12). Milk-bottle opening is of particular interest because it represents one of the best-known examples of non-human animal *culture*. Culture is thought to represent an additional form of inheritance that fits into a wider framework alongside genetic inheritance, the epigenome and ecological factors such as territories (13,14). These innovations can result in behavioural traditions when they spread through a population via social learning *and* persist over time, and it is the sum of these traditions that constitutes a population’s culture (15). The benefits of this second form of inheritance are immediately apparent: while genetic inheritance acts solely on a vertical basis, cultural transmission occurs both vertically and horizontally (16). This allows a more rapid response to changes in the environment than genetic transmission alone. Culture may also influence genetic inheritance to the extent of driving speciation: behavioural differences between orca ecotypes that exist in sympatry has been argued to result in reproductive isolation so complete that, if functional speciation has not already occurred, it will inevitably do so (17–19). Another potential mechanism through which cultural and genetic inheritance might intersect is the phenotype-first theory of evolution, otherwise known as the Baldwin effect. Here, beneficial behavioural traits acquired during life are passed on to offspring via selection that favours the acquisition of such behaviour, such as on learning ability or behavioural biases (20,21). If a learned, beneficial behavioural innovation were to be maintained in a population over multiple generations by social learning, it seems likely that selection might act to favour variants that are more capable at learning the behaviour in question.

The idea that what now appears merely instinctive may have originated via learning, or even culture, has the potential to explain the evolutionary origin of many complex behaviours. We here explore the possibility that cultural processes might, at least in theory, have contributed to the advent of unique behavioural innovations in social insects, using bumblebees. In the laboratory, bumblebees have been shown to acquire both simple information such as flower colour choice (22,23) and relatively complex, non-natural foraging techniques from other bees, such as string-pulling, in paired dyad paradigms (24). However, the spread of such techniques has not yet been observed under the conditions that are considered the gold standard for demonstrating culture in the laboratory: the so-called open diffusion paradigm (25). These experiments are of high ecological validity and involve the release of a trained demonstrator into a group of naïve observers, along with provision of the substrates necessary to perform the target behaviour.

Cultural variation serves as the raw substrate for cultural evolution, and it is for this reason that investigating the capacity for cultural variation is as important as investigating the capacity for culture itself. At minimum, to investigate cultural variation, a two-action control design is required. One such study involved training great tit demonstrators to open a puzzle box in one of two possible ways before seeding them back into wild populations (26). The demonstrator’s preferences spread throughout these groups and were maintained long-term, even when the alternative behaviour was discovered, and even though the two variants were entirely arbitrary. Thus, to investigate the capacity for culture and its dynamics in bumblebees, we designed two-option puzzle box feeders informed by previous work on bumblebee problem-solving (27), that replicated those used to investigate arbitrary traditions in great tits. We then developed an open diffusion protocol that allowed the spread of box-opening to be recorded, and seeded colonies of bumblebees with demonstrators trained to perform one of the two possible behavioural variants. These novel foraging techniques were acquired by untrained bees via social learning, and the maintenance of local variations of this behaviour indicated a capacity for culture in bumblebees.

## Results

### Box-opening behaviour spread through bumblebee colonies under open diffusion conditions

To determine whether bumblebees could acquire and sustain cultural variation, we designed two-option puzzle boxes that could be opened by rotating a clear lid around a central axis by either pushing a red tab clockwise or a blue tab counter-clockwise (termed the “red-pushing behavioural variant” and “the blue-pushing behavioural variant” respectively) to expose a 50% sucrose solution reward, as indicated by a yellow target (Fig. 1A). Video files showing bees performing both behavioural variants are available in the *Supporting information*. Stages of the incremental training protocol developed for demonstrators are depicted in Fig. 1B, and full details of this and the open diffusion protocol can be found in the *Materials and Methods*. Demonstrators trained to perform the red- and blue-pushing behavioural variants will henceforth be referred to as “red-pushing demonstrators” and “blue-pushing demonstrators”, respectively. We conducted three experiments in total, with *Experiment 1* and *2* involving the seeding of a single trained demonstrator into a population (and open diffusion experiments being conducted over 6 or 12 consecutive days, respectively). We also provided opportunities for bees to innovate and solve the boxes without social input, in control populations where no demonstrator was present. *Experiment 3* involved the seeding of multiple demonstrators into a population (including two red-pushing demonstrators and two blue-pushing demonstrators) and was conducted over 12 consecutive days.

**Figure 1.**
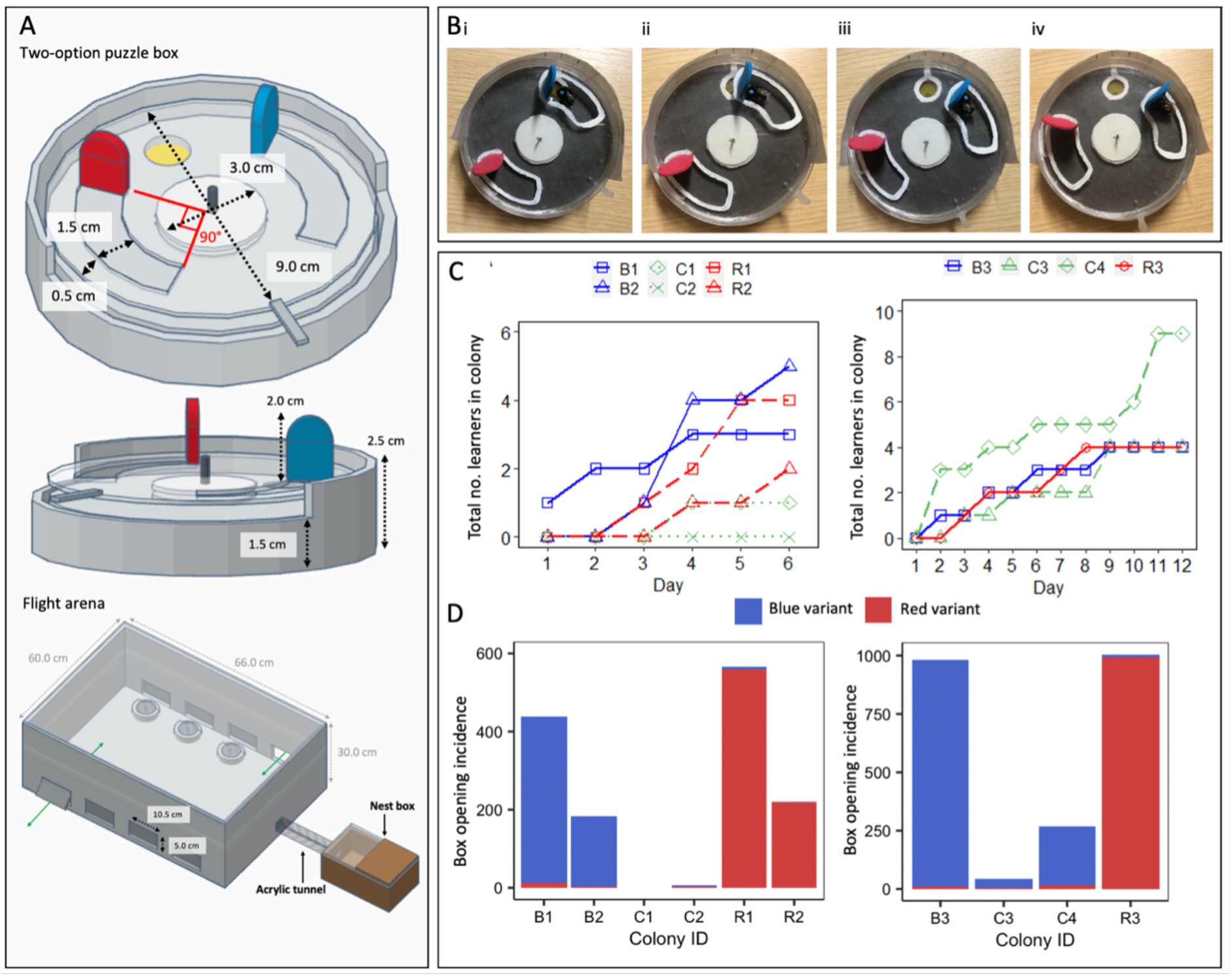
Box-opening behaviour spreads through bumblebee populations under open diffusion conditions. **(A) Diagrams of puzzle boxes and the experimental flight arena used for the single-demonstrator diffusion experiments.** Boxes were constructed from Petri dishes, with a base and lid serving as the static lower part of the box. This was covered with laminated grey, RGB neutral paper (hex #555555), which was lightly sanded to provide grip. An additional Petri dish lid with two cut-out tracks served as the rotating lid of the box. White acrylic “washers” and a nail formed the rotating mechanism. Red and blue tabs were cut from 2 mm-thick polyethylene craft foam sheets and affixed to one end of the cut-out tracks as depicted; thus, when a bee landed in either track and pushed against its tab, walking forwards, it would rotate the lid of the puzzle box around the central axis and expose the reward (50% w/w sucrose solution on the yellow target). An external plastic “shield” encircled the box to prevent bees obtaining the reward by squeezing in at the sides, and a stopper prevented the lid from being over-rotated after the reward was exposed. The flight arena was connected to the nest box via an acrylic tunnel, and flaps cut in the side of the flight arena allowed the removal and replacement of puzzle boxes during the experiment. The sides were lined with bristles to prevent bees escaping. **(B) Incremental demonstrator training protocol.** Panels show the procedure to train a bee to perform the anti-clockwise “blue-pushing behavioural variant”, to train a demonstrator to perform the clockwise “red-pushing behavioural variant”, the red tab would start over the reward instead. (i) Initial configuration with the box fully open and the yellow target completely exposed and accessible. (ii) In this configuration, the reward can still be obtained by reaching under the tab with the proboscis, but the tab is often pushed forward as the bee attempts this. (iii) The reward can no longer be obtained without moving the tab forwards. (iv) The two tabs are almost equidistant, with the blue tab ∼1.0cm closer to the yellow target than the red. Once a bee solved a box with this configuration twice in a single foraging bout, it progressed to the unrewarded learning test. **(C) Diffusion curves for Experiment 1 (left panel) and Experiment 2 (right panel).** Observers were considered to have learned after opening a box twice; they were only defined as having opened a box whenever they pushed either tab ≥50% of the required distance, obtaining the reward. **(D) Overall box opening incidence by learners in Experiment 1 (left panel) and Experiment 2 (right panel).** The incidence of each behavioural variant is indicated by colour. Colonies B1, B2 and B3 were each seeded with a blue-pushing demonstrator, colonies R1, R2 and R3 were each seeded with a red-pushing demonstrator, and colonies C1, C2, C3 and C4 were controls that lacked a demonstrator. Data for the whole colonies, including the demonstrator’s behaviour, can be found in Appendix Fig. 1-2 and Appendix Table I.

For the open diffusion experiments, eight puzzle boxes were presented in the flight arena and all bees were allowed into the flight arena to freely interact with these boxes. Each day, bees received 30 min pre-training with lidless boxes, which left the yellow target fully accessible and allowed bees to form an association between the yellow target and the 50% w/w sucrose solution reward. This training period was followed by a 3 h experimental period, during which the puzzle boxes were closed and bees had to push on one of the two tabs to access the reward. Once each box had been opened and the sucrose reward consumed, it was removed from the arena. During this period, we monitored how often each box was opened and determined the identity of the bee responsible for each opening.

A total of six bumblebee colonies were used for the 6-day, single-demonstrator open diffusions of *Experiment 1* (two colonies seeded with a red-pushing demonstrator and two seeded with a blue-pushing demonstrator, and two control colonies). Untrained bees opened puzzle boxes in all four experimental colonies and a number of these bees (n=14) met the learning criterion as defined in the *Materials and Methods* (Fig. 1C). In contrast, in the two control colonies, only one individual opened a box, despite these colonies being of a comparable size to the experimental colonies. This bee did proceed to meet the learning criterion (Fig. 1C), although it only opened boxes sporadically (n=5 incidences in total; Fig. 1D).

### Bees that met the learning criterion in the presence of a trained demonstrator were significantly more proficient than those that did so without

The emergence of this spontaneous learner in the absence of any demonstrator necessitated *Experiment 2*: single-demonstrator open diffusions run for 12 days instead of 6, to determine whether more spontaneous learners would emerge over a longer period. A total of four colonies were used for these experiments (two experimental colonies, with one seeded with a red-pushing demonstrator and the other with a blue-pushing demonstrator, and two control colonies). Untrained bees from all four colonies, experimental and control, reached the criteria to be considered learners (Table 1). Remarkably, the highest number of individuals met learning criteria in colony C4 (n=9 vs. n=4 in each experimental colony).

Although box opening could evidently arise spontaneously in the absence of social learning, we found that bees that learned from demonstrators were significantly more proficient at box opening than learners from control colonies (Fig. 1D). We pooled learners from *Experiments 1* and *2* and calculated individual proficiency indices to allow comparison between bees who met the learning threshold at different points in the diffusion (and thus had more or less chance to accumulate box openings). These indices were calculated as follows: total individual opening incidence / total days spent as a learner, inclusive of the day the learning threshold was met. The difference between learners from experimental and control colonies was significant, with learners from experimental colonies opening significantly more boxes than spontaneous learners from control colonies (median boxes opened per day after reaching learning threshold: experimental colonies, 27.9 (IQR=2.25-60.6); control colonies, 1.15 (IQR=0.69-3.42); Mann-Whitney U-test: W=253, p=0.001; Fig. 2A). This individual-level difference in proficiency translated into colony-level differences in the frequency of box opening (median boxes opened per day by learners, experimental colonies: 76.8 (IQR=45.4-83.2); control colonies, 2.1 (IQR=0.6-8.1); Mann-Whitney U-test: W=28, p=0.006). This suggested that, although social learning was not required for a bee to perform box-opening behaviour, it was necessary for the behaviour to be repeated frequently: i.e. to become fixed in an individual’s (or population’s) repertoire.

**Table 1.** Number of untrained bees to open the puzzle box and meet learning criteria in the single-demonstrator diffusion experiments.

| Colony ID<sup>1</sup> | Diffusion length (days) | No. observers to open a box | No. observers to meet learning criterion<sup>2</sup> | Box opening incidence by all observers | Box opening incidence by learners |
| --- | --- | --- | --- | --- | --- |
| B1 | 6 | 8 | 3 | 437 | 432 |
| B2 | 6 | 6 | 5 | 182 | 181 |
| B3 | 12 | 5 | 4 | 980 | 979 |
| R1 | 6 | 4 | 4 | 565 | 565 |
| R2 | 6 | 2 | 2 | 219 | 219 |
| R3 | 12 | 5 | 4 | 1006 | 1005 |
| C1 | 6 | 0 | 0 | 0 | 0 |
| C2 | 6 | 1 | 1 | 5 | 5 |
| C3 | 12 | 4 | 4 | 41 | 41 |
| C4 | 12 | 14 | 9 | 269 | 264 |

<sup>1</sup>Colonies B1-3 were seeded with a blue-pushing demonstrator, colonies R1-3 with a red-pushing demonstrator, and colonies C1-4 were controls that lacked a demonstrator. <sup>2</sup>Observers were considered to have learned after opening a box twice; they were only defined as having opened a box whenever they pushed either tab $\geq 50\%$ of the required distance, obtaining the reward (see Materials and Methods for full detail).

**Figure 2.**
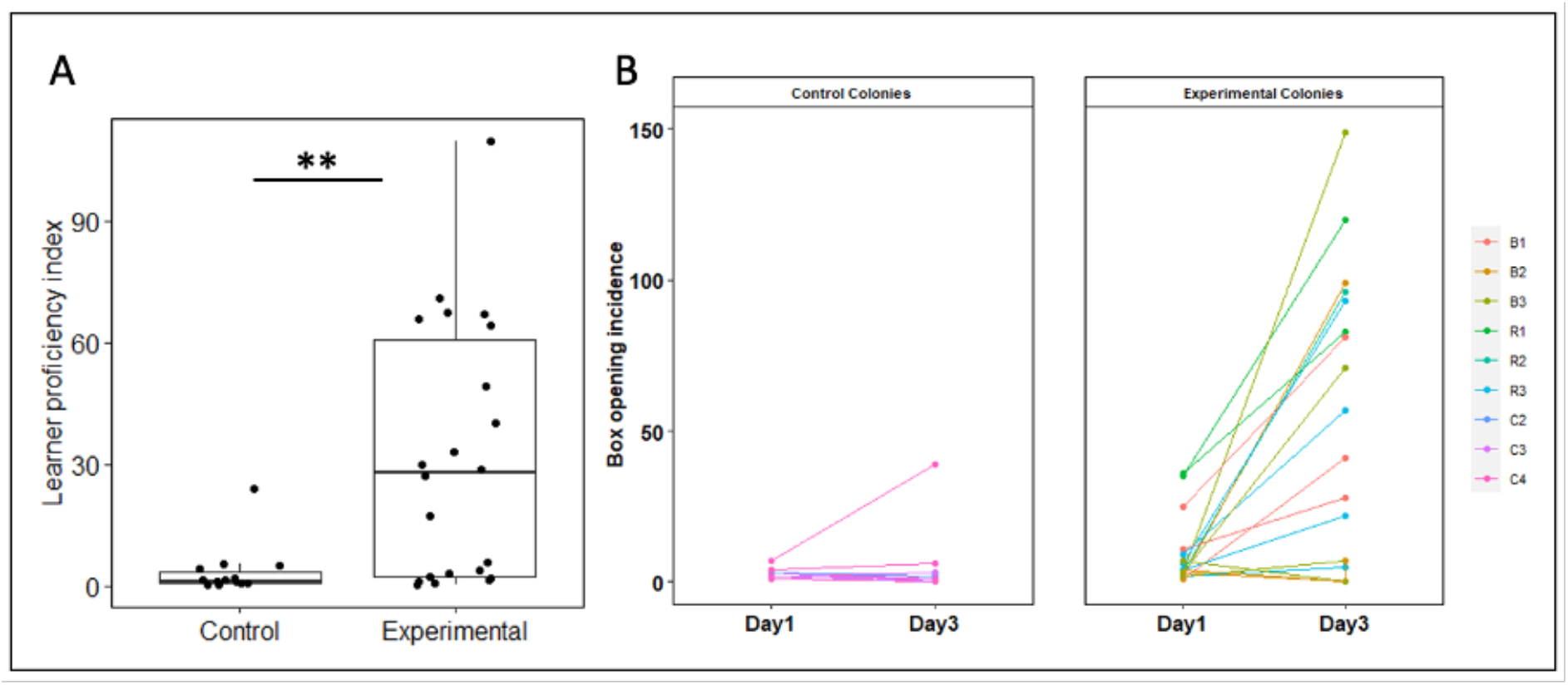
Learner proficiency is higher in bees exposed to a demonstrator compared with those without, and increases more rapidly. **(A) Difference in proficiency between learners from experimental and control colonies.** Proficiency indexes were calculated for each individual learner as follows: total incidence of box opening / days spent as a learner, with the latter spanning from the day learning criteria was met to the end of the diffusion. Control, n=14, including data from colonies C2-4; experimental, n=22, including data from colonies R1-3 and B1-3.) **P<0.01. (B) **Change in learner proficiency from the day learning criteria were met (“day 1”) to two days later (“day 3”) in control and experimental diffusion experiments.** Learners who did not meet the learning criteria early enough to have data for a third day (e.g. those who met criteria on day 5 or 6 of the 6-day colonies or on day 11 or 12 of the 12 day colonies) were excluded from analysis. Control group, n=11; experimental group, n=18.

Next, we sought to analyse the effect of demonstrator presence on learner proficiency over time. To achieve this, we compared the incidence of box openings on the day an individual met learning criteria (termed ‘day 1’) and two days later (termed ‘day 3’; Fig. 2B). Analysis with a linear mixed-effects model identified a statistically significant interaction between treatment (exp vs. ctrl) and day (day 1 vs. day 3), with bees in the experimental group experiencing a greater increase in proficiency on day 3 compared with day 1 than bees in the control group (F=10.152, df=1, p=0.003; Appendix Table 2). This confirmed that the presence of a demonstrator led to increased proficiency and more persistent behaviour amongst learners in the experimental colonies, indicating that social learning was crucial to the long-term persistence of novel behaviours and behavioural variants within the population.

**Table 2.**
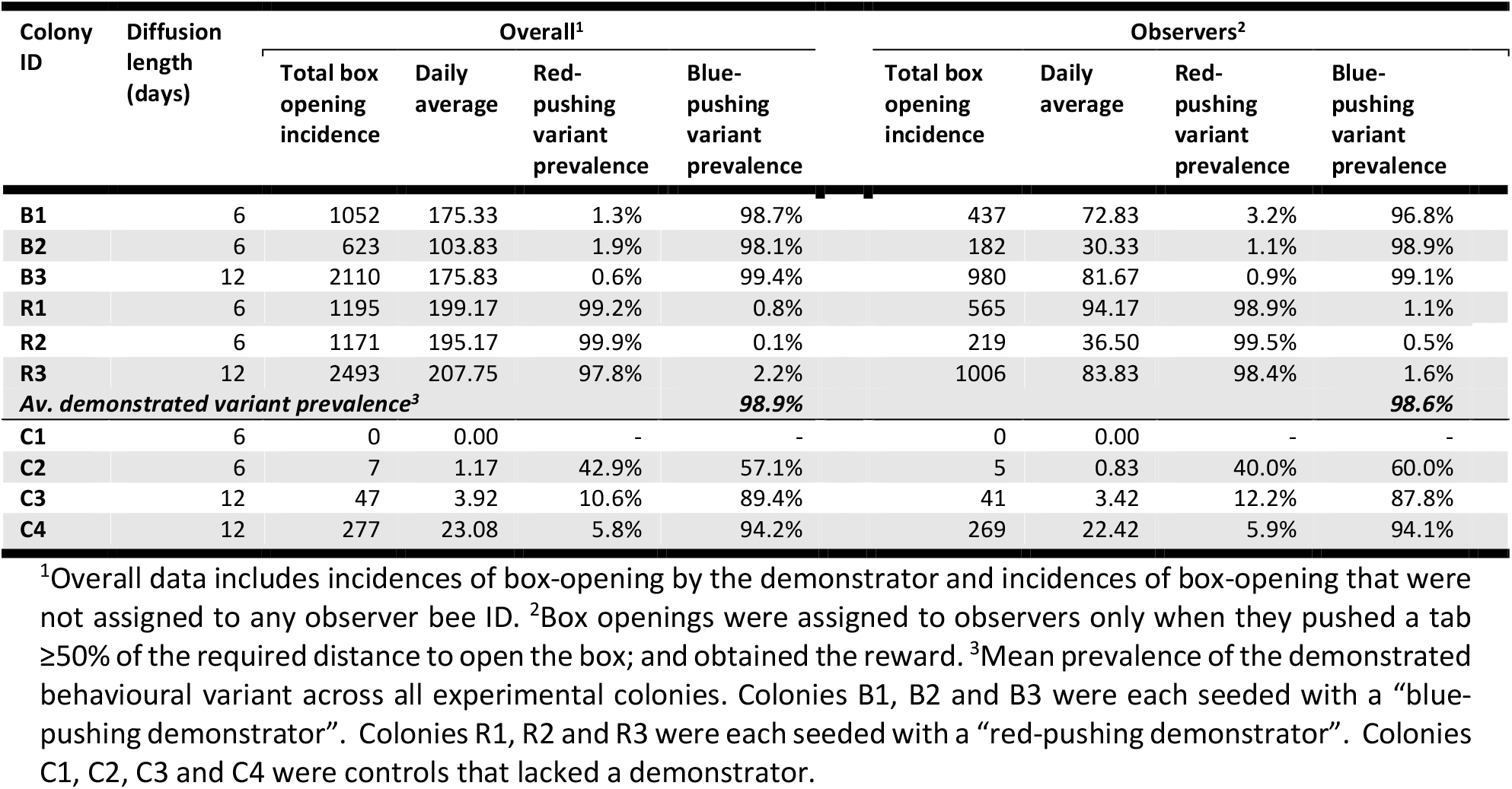
Preferences for puzzle box-opening behavioural variants in the single-demonstrator diffusion experiments.

### When exposed to demonstration of a single arbitrary behavioural variant, learners acquired a strong preference for this variant that persisted even when the alternative behaviour was discovered

Puzzle boxes could be opened in one of two ways: by pushing either the red tab clockwise, or the blue tab anticlockwise. In every experimental colony, the behavioural variant performed by the demonstrator became dominant among the learners (Table 2). This preference remained even when learners discovered the alternative behaviour: more than half the learners from the experimental colonies performed the non-demonstrated variant at least once (12 out of 22 bees; Appendix Tables 3 and 4). Nonetheless, individual learners from experimental colonies were significantly more likely to perform their taught variant than the alternative (median proportion of box-openings made using the taught variant, 1.0 (IQR=1.00-0.99; taught variant incidence vs. alternative variant incidence: Wilcoxon signed-rank test, V=531, p<0.001). There was no difference between the strength of preference for the demonstrated variant between individual learners from colonies seeded with blue- or red-pushing demonstrators (Mann Whitney U test, W=95, p=0.279). This individual-level preference for the taught variant translated into a striking colony-level trend, with a mean of 98.6% of box openings made using the taught variant. Even in colony B1, where the preference for the demonstrated variant was weakest, it was still performed by observers 96.8% of the time (Table 2).

**Table 3.** Learner characteristics in the multiple-demonstrator diffusion experiments.

| Bee ID | Colony ID | Day learning criteria met | Incidence of box-opening behaviour |  |  |  |  |
| --- | --- | --- | --- | --- | --- | --- | --- |
|  |  |  | Total | Red variant | Blue variant | % Red variant | % Blue variant |
| Proficient learners |  |  |  |  |  |  |  |
| g58 | 1R2B2 | 2 | 962 | 34 | 928 | 3.5 | 96.5** |
| b18 | 1R2B2 | 2 | 1664 | 1627 | 37 | 97.8** | 2.2 |
| r63 | 1R2B2 | 6 | 44 | 21 | 23 | 47.7 | 52.3 |
| g12 | 1R2B2 | 5 | 471 | 428 | 43 | 90.9** | 9.1 |
| w75 | 1R2B2 | 5 | 99 | 58 | 41 | 58.6 | 41.4 |
| w50 | 1R2B2 | 6 | 44 | 27 | 17 | 61.4* | 38.6 |
| g49 | 1R2B2 | 8 | 268 | 2 | 266 | 0.7 | 99.3** |
| g50 | 1R2B2 | 8 | 11 | 3 | 8 | 27.3 | 72.7* |
| y70 | 1R2B2 | 8 | 19 | 16 | 3 | 84.2** | 15.8 |
| Av. |  |  | 398.0 | 246.2 | 151.8 |  |  |
| y47 | 2R2B2 | 1 | 24 | 19 | 5 | 79.2* | 20.8 |
| g56 | 2R2B2 | 2 | 832 | 12 | 820 | 1.4 | 98.6** |
| g24 | 2R2B2 | 4 | 388 | 10 | 378 | 2.6 | 97.4** |
| r50 | 2R2B2 | 8 | 12 | 3 | 9 | 25.0 | 75.0* |
| Av. |  |  | 314.0 | 11.0 | 303.0 |  |  |
| Non-proficient learners |  |  |  |  |  |  |  |
| y1 | 1R2B2 | 2 | 2 | 1 | 1 | 50.0 | 50.0 |
| b54 | 1R2B2 | 9 | 2 | 1 | 1 | 50.0 | 50.0 |
| g79 | 1R2B2 | 3 | 9 | 0 | 9 | 0.0 | 100.0 |
| g84 | 1R2B2 | 4 | 2 | 2 | 0 | 100.0 | 0.0 |
| r59 | 1R2B2 | 7 | 2 | 1 | 1 | 50.0 | 50.0 |
| g3 | 1R2B2 | 8 | 2 | 1 | 1 | 50.0 | 50.0 |
| g54 | 1R2B2 | 8 | 4 | 0 | 4 | 0.0 | 100.0 |
| g44 | 1R2B2 | 12 | 2 | 1 | 1 | 50.0 | 50.0 |
| r52 | 1R2B2 | 12 | 3 | 3 | 0 | 100.0 | 0.0 |
|  |  | Av. | 3.1 | 1.1 | 2.0 |  |  |
| g54 | 2R2B2 | 3 | 2 | 0 | 2 | 0.0 | 100.0 |
| y20 | 2R2B2 | 4 | 3 | 1 | 2 | 33.3 | 66.7 |
| y55 | 2R2B2 | 5 | 7 | 0 | 7 | 0.0 | 100.0 |
| w64 | 2R2B2 | 12 | 2 | 1 | 1 | 50.0 | 50.0 |
|  |  | Av. | 3.5 | 0.5 | 3.0 |  |  |

Asterisks represent the variant preference (following the figure for the preferred variant) and the strength of preference (\*\* indicating a strong preference, and \* a weak preference). As the sample sizes for non-proficient learners were small, no preferences or strength of preferences were assigned.

Learners from control colonies had no taught variant but preferred to open boxes using the blue-pushing variant (median proportion of box-openings made using the blue-pushing variant, 0.99 (IQR=1.00-0.78). While this preference was not as strong at the colony level as in the experimental colonies (C2, 60.0%; C3, 87.8% and C4, 94.1% prevalence; Table 2), it did suggest an innate inclination to perform the blue-pushing behavioural variant, an unsurprising result in light of the well-known preference for the colour blue among bumblebees (28,29). There was also no significant difference between the strength of preference between control and experimental colonies: control learners preferred their favoured variant just as strongly as experimental learners (Mann Whitney U test, W=379, p=0.164). The fact that learners exposed to a red-pushing demonstrator showed just as strong a preference for the red-pushing variant as control learners did for their preferred variant, even in the face of a natural inclination towards the blue-pushing variant, provides further evidence suggesting that social learning is key to the transmission of puzzle-box opening. In short, a learned preference for red can overcome an innate preference for blue.

Overall, the results of Experiment 1 and 2 suggest that spontaneous learners from control colonies are less proficient at box-opening than social learners, opening fewer boxes in total. Social learners acquired strong preferences for the demonstrated variant even though more than half tried the alternative variant. This translated to greater proficiency and strong preferences for a single variant at the colony level.

### Box-opening behaviour persisted among learners in experimental colonies but was vulnerable to collapse in the controls

Experiment 2 also provided some insight as to the likelihood of box-opening behaviour persisting over time and/or generations of learners; a prerequisite for it to be considered cultural in nature. Figure 3 depicts the daily incidence of box-opening behaviour by observers throughout all single-demonstrator diffusions (full data is presented in Appendix Fig. 1 and Appendix Table 1). The incidence of box-opening by observers was significantly positively correlated with experimental day in 5/6 experimental colonies, while there was no significant correlation in 3/4 control colonies (Spearman’s rank order correlation tests; Fig. 3, Appendix Table 1). The control colonies also appeared to be subjected to greater fluctuation: while peaks and troughs in incidence do occur in the experimental colonies, box-pushing still occurred relatively frequently on these days compared with control colonies. In colony C3, box-opening appears to arise on two separate occasions (day 3 and day 9), separated by a complete collapse on day 6. Our diffusion experiments were run for 12 days at the maximum and did not include defined generations of learners, but the results do indicate that behaviours arising in control colonies are less likely to persist over time compared with those in the experimental colonies, even when the number of learners is comparable. Taken together, these results suggest that it was social learning that underpinned the spread and maintenance of box-opening and its variants in the experimental colonies under open diffusion conditions.

**Figure 3.**
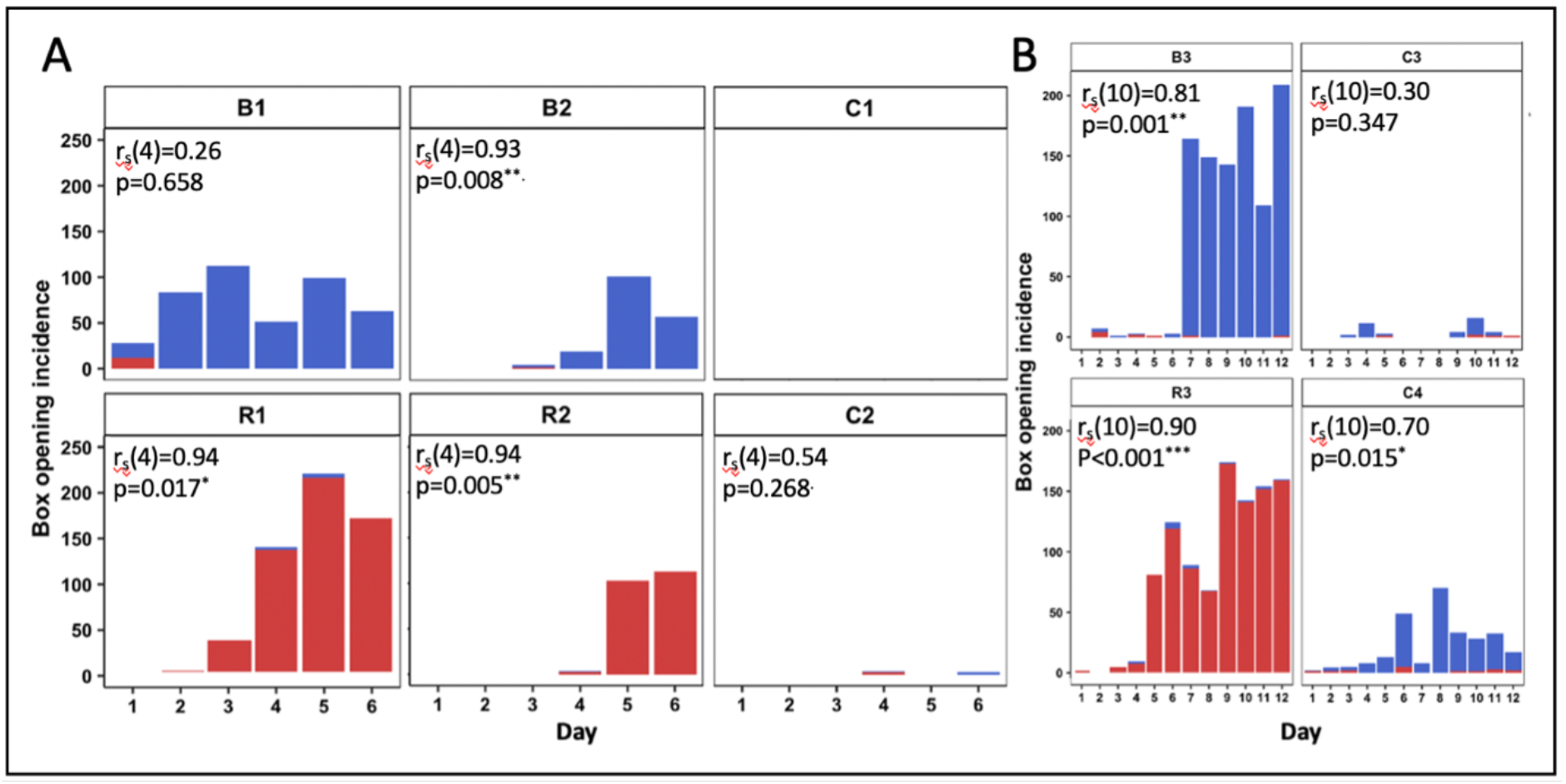
Daily incidence of box openings by observers in (A) Experiment 1 and (B) Experiment 2. Colonies R1, R2 and R3, and B1. B2 and B3 were seeded with single demonstrators trained to perform the red-pushing behavioural variant and the blue-pushing behavioural variant, respectively. Colonies C1, C2, C3 and C4 were controls that lacked a demonstrator. A box opening was assigned to an observer whenever it pushed on either tab ≥50% of the required distance and obtained the reward. Incidences of the “blue-pushing behavioural variant” are depicted in blue, while incidences of the “red-pushing behavioural variant” are depicted in red. Data for the whole colonies and demonstrator activity can be found in Appendix Fig. 1 and Appendix Table 1. Relationships between experimental day and opening incidence (of any variant) by observers were analysed using Spearman’s rank order correlation tests. ^*^p<0.05, ^**^p<0.01 and ^***^p<0.001.

### When exposed to both behavioural variants in roughly equivalent proportion, one behavioural variant became dominant in each population

The experiments described in this article thus far (and, indeed, most open diffusion experiments conducted in the literature) begin with arbitrary local variation already established (i.e., demonstrators are trained to perform a single behavioural variant ∼100% of the time and the alternative is left to be discovered serendipitously, if at all). Such experiments are usually conducted to see whether fidelity to this tradition would degrade over time or be sustained. Less work, comparatively, has been done on *how* such a local behavioural variation might emerge in the first place within a population. Thus, we conducted an additional experiment aimed at investigating what might happen if two behavioural variants were initially present in a population. *Experiment 3* involved the seeding of multiple demonstrators into a population, and used the same puzzle boxes and training protocol as *Experiment 1* and *2*. However, the flight arena was modified to permit the presentation of 16 puzzle boxes and the simultaneous exposure of two colonies, providing a larger population of foragers (Fig. 4A). A total of four demonstrators were seeded into the population, with two trained to perform the red-pushing behavioural variant and two the blue, and the diffusion was conducted for 12 days. This experiment was run twice, with the replicates termed population *1R2B2* and *2R2B2*. In each replicate, demonstrators opened boxes using the two behavioural variants in close to equal proportions (proportion of all openings made with blue variant: 1R2B2, 0.550; 2R2B2, 0.564; Appendix Fig. 3; Appendix Table 5).

**Figure 4.**
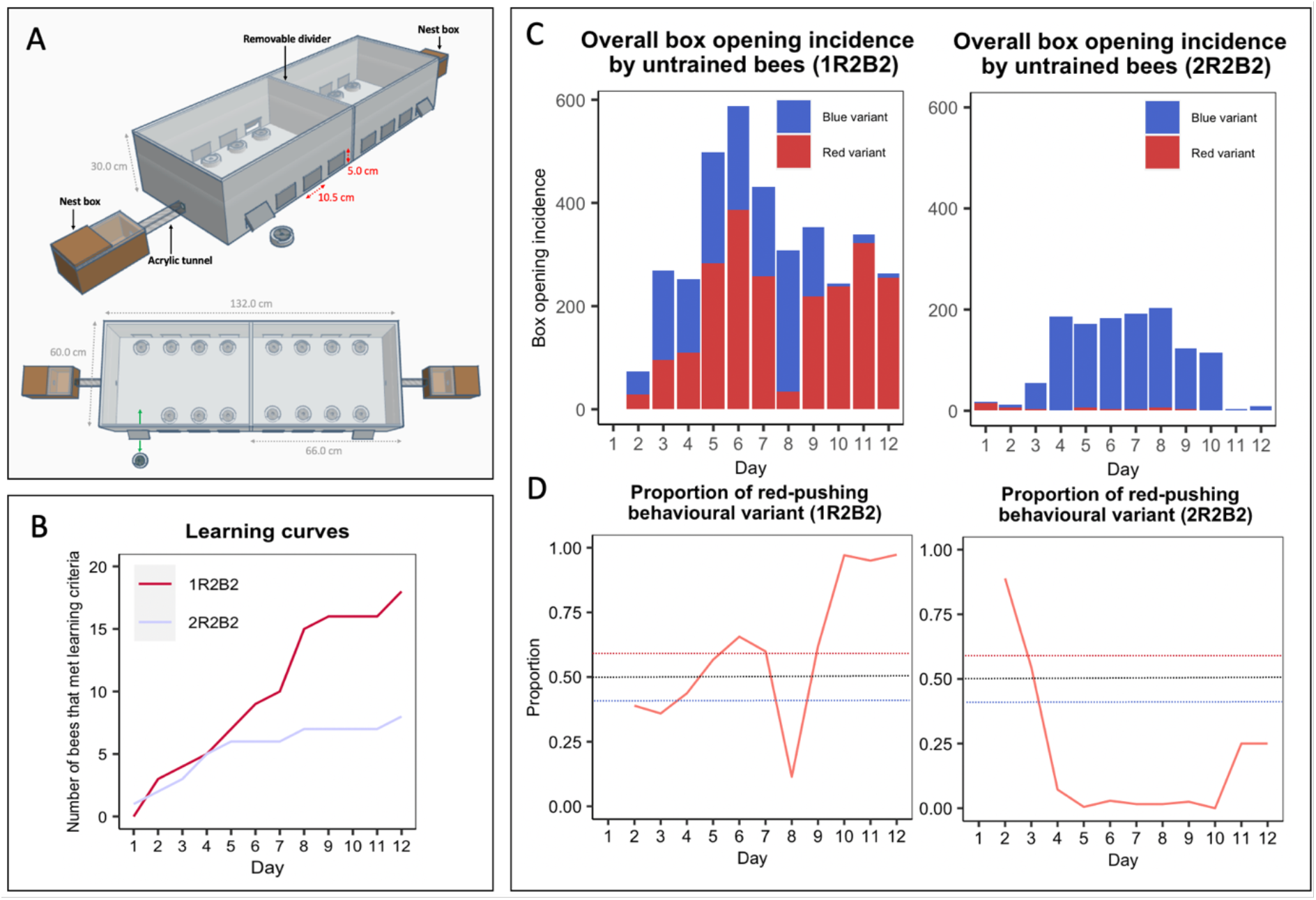
When exposed to simultaneous demonstration of both behavioural variants, one variant becomes dominant among learners. **(A) Multiple-demonstrator puzzle box open diffusion experimental set-up.** The multiple-demonstrator diffusion flight arena was bipartite, with a removable divider splitting the arena in two, and allowing 16 boxes to be presented simultaneously. Flaps cut in the side of the flight arena allowed the removal and replacement of puzzle boxes during the experiment, and the sides were lined with bristles to prevent bees escaping. **(B) Diffusion curves for Experiment 3.** Observers were considered to have learned after opening a box twice; they were only defined as having opened a box whenever they pushed either tab ≥50% of the required distance, obtaining the reward. **(C) Overall box opening incidence by learners in Experiment 3 (left panel, population 1R2B2, right panel, population 2R2B2).** The incidence of each behavioural variant is indicated by colour. **(D) The proportion of recorded daily behaviours by learners that were the red-pushing variant.** Dashed lines show the thresholds for a preference for either variant. Overall data, including incidences of box-opening by the demonstrator and incidences of box-opening that were not assigned to any observer bee ID, are presented in Appendix Fig. 4. Full dataset is available in Appendix Table VI.

A total of 18 observers met the learning threshold in population 1R2B2, with 9 of these performing box-opening behaviour >10 times and thus being termed *proficient learners*. In 2R2B2, 8 observers met the learning criterion, and 4 of these were classed as proficient learners (Fig. 4B, Table 3, Appendix Table 7). Notably, by day 12 of diffusion, the red-pushing behavioural variant became the dominant technique among observers in population 1R2B2, while the blue-pushing behavioural variant predominated in population 2R2B2 (Fig. 4C and D; Appendix Table 6). Strikingly, all demonstrators in this population had ceased their activity by day 6; all box-opening during the remaining six days was performed by observers alone (Appendix Table 5). Four new individuals reached the learning threshold after day six in this colony, suggesting that they may have learned from the previous generation of learners rather than the demonstrators (Table 3).

In population 1R2B2, until day 8, the predominance of the red-pushing and blue-pushing behavioural variants fluctuated back and forth somewhat evenly (Fig. 4C and D). However, from day 9 onwards, the red-pushing behavioural variant became increasingly predominant, and by day 12, 97.3% of the 263 incidences of observer box-opening behaviour were of the red-pushing variant. In population 2R2B2, observers preferred the blue-pushing variant over the red on all days except day 1, where 16/18 box-opening incidences by observers were of the red variant. However, on days 11 and 12, there was a sharp drop in observer activity (Fig. 4C, Appendix Table 6), despite no apparent fall in box-opening overall (Appendix Fig. 3A, Appendix Table 6). This was driven by the activity of two trained demonstrators, which both remained active on day 12.

### When faced with demonstrations of both variants, proficient learners tended to form preferences for a single variant

Proficient learners, who performed box-opening behaviour >10 times, formed strong individual preferences for a single variant with few exceptions (median proportion of box-openings made using the preferred variant, 0.84 (IQR=0.97-0.73; preferred variant incidence vs. non-preferred variant incidence: Wilcoxon signed-rank test, V=91, p<0.001 vs. chance level). Notably, in both populations, some individuals developed a preference for the blue-pushing variant, and others for the red-pushing variant. Even though learners generally performed both behavioural variants during the experiment, once a learner developed a preference for either variant, they maintained this preference and showed no evidence of switching variants (Fig. 5): there was no significant difference between in variant preference between the first day of learning and the last day of recorded activity (linear mixed effects model, F=0.626, df=1, p=0.4377; Appendix Table 8). In one case (b18), a strong preference for the red behavioural variant persisted even after a two-day pause in foraging activity (Fig. 5).

**Figure 5.**
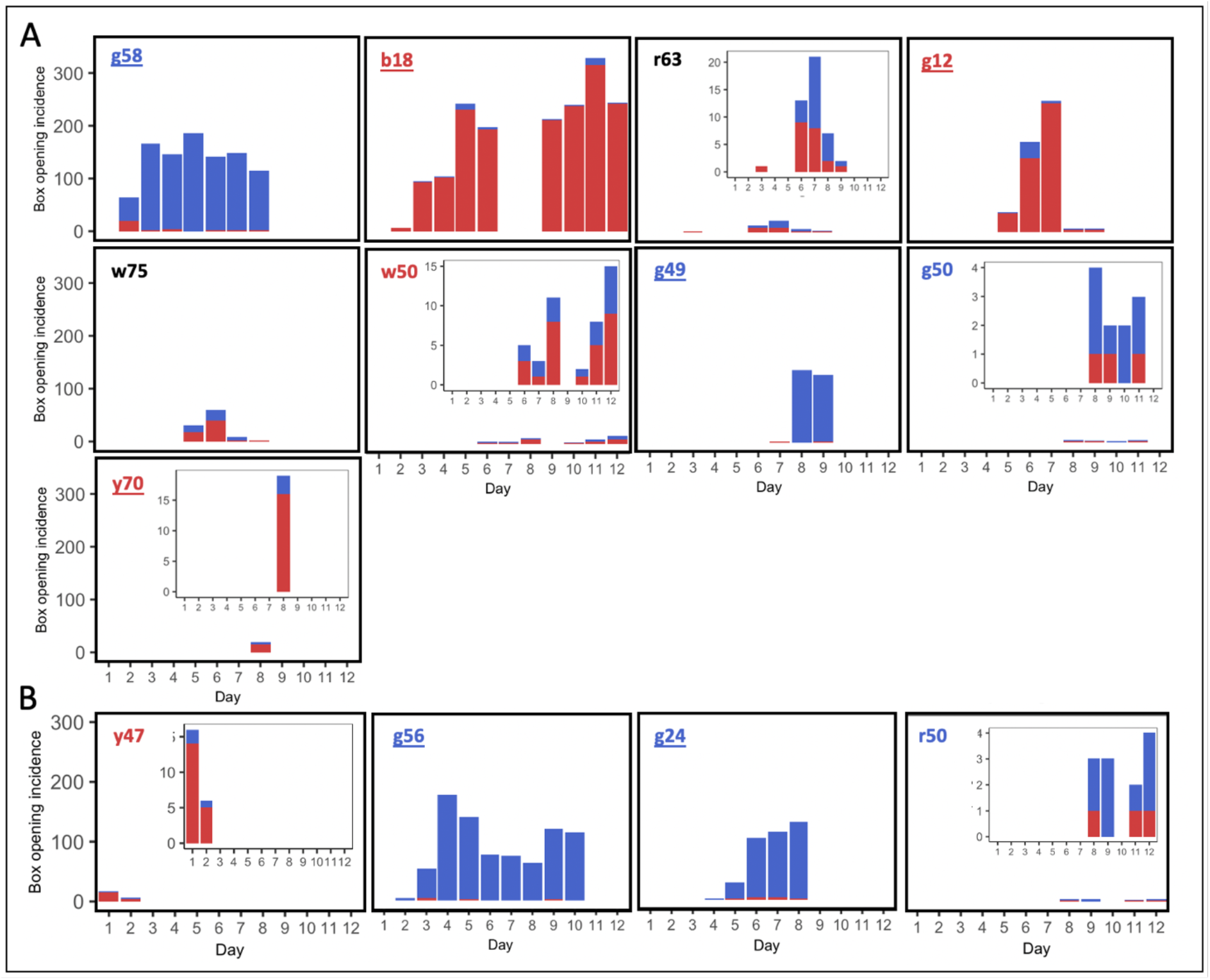
Box opening incidence for individual proficient learners from (A) colony 1R2B2 and (B) 2R2B2. Overall preferences are indicated by the colour of the graph titles, and strong preferences are indicated by an underline. Inset graphs show the same data at a finer scale, in cases where daily incidence is especially low and so difficult to read on the common scale.

Shifts in colony-level preference appeared to be due to the emergence of new learners (that developed their own individual preference) and the cessation of foraging activity by individuals with established preferences, rather than currently active foragers changing their behaviour to match the current majority.

## Discussion

The capacity for culture was initially thought to be uniquely human, but behaviours that meet the prerequisites to be considered traditions have since been found to occur naturally in a number of animals. These include tool selection in chimpanzees (30), song dialects in birds (31,32), and feeding techniques in humpback whales (33). However, the majority of these studies focus on vertebrates with relatively large brains.

The results of the present study provide strong evidence that social learning underpins the transmission of novel foraging behaviours in bumblebees, but most importantly that these behaviours (and arbitrary variants of these) can spread and persist under open diffusion conditions. This fulfils the requirements for these behaviours to be considered traditions: they must be acquired socially and maintained by a population over time (15). Notably, in the experimental single-demonstrator diffusions, some untrained observers discovered the alternative, non-demonstrated technique but reverted to the demonstrated variant. This suggests that although bees were able to generalise their behaviour and were not entirely ignorant of the alternative technique, they still preferred the variant that was demonstrated to them. This result matches those found in great tits, which also discovered the alternative technique for acquiring a food reward from a puzzle box, but then reverted to the demonstrator’s preferred behaviour (26). Furthermore, the results of the multiple-demonstrator diffusion experiments suggest that when more than one behavioural variant is present in a bumblebee population, arbitrary local traditions may emerge over time. To our knowledge, this is the first demonstration of the rise, spread and maintenance of arbitrary behavioural variation under open diffusion conditions in any invertebrate.

But how can we claim with confidence that social learning was occurring when some individuals in control diffusion experiments opened puzzle boxes without a demonstrator? This behavioural flexibility, remarkable in itself, does not undermine the conclusion that box-opening behaviour represents a behavioural tradition in bumblebees. For one thing, all culture requires the capacity for behavioural innovation in the first place, so its existence alone cannot act as evidence against culture. Furthermore, the proficiency of these control learners was lower compared with those from the experimental diffusions, despite these learners needing to compete with other (more proficient) learners *and* a highly proficient trained demonstrator for boxes. If bees from the control diffusions had learned in the same way as those in experimental diffusions, they should have been able to accumulate more openings due to lack of competition. Furthermore, bees in the control diffusions should have been more driven to forage: for the full three hours of the daily open diffusion, the experimental colonies enjoyed an influx of high-value 50% w/w sucrose solution from successful foragers while the control colonies received little, or none.

The daily pretraining ensured that bees in both conditions knew that the yellow target on the puzzle box indicated a reward. When they could no longer access this during the open diffusion, but could still see the yellow target through the lid, some bees in the control colonies began to investigate the target and other components of the box. Eventually, this would inadvertently move either the blue or red tab closer to the target (see media file 3). If this behaviour was undirected or very incremental, perhaps it was simply not possible for bees to form an association between tab-pushing and it with the reward. In contrast, bees in the experimental colonies had their attention repeatedly drawn towards relevant features of the box via the demonstrator, and thus local enhancement. This may have resulted in their own attempts to open boxes being more directed. Alternatively, the constant, repeated reinforcement of the tab being juxtaposed with the target, as in opened boxes, may have permitted the association to form via stimulus enhancement. Bees in the control colonies that met learning criteria may well have experienced some degree of social input: once one bee opened a box, it might have acted as a demonstrator for others. But if these initial learners and those that followed them were not proficient, subsequent learners may not have received enough social input. The trained demonstrators were highly proficient, and regularly opened boxes more than 100 times a day (Appendix Table I). In contrast, the most proficient control learner, bee y12 from colony C4, opened boxes just 68 times on its most successful day (Appendix Table V). This particular bee was an outlier with 216 box-openings recorded during the experiment (for scale, its closest rival among the control learners opened boxes just 22 times in total; Appendix Table V). From this, it seems clear that it was the presence of the demonstrator that caused box-opening behaviour to become fixed in an observer’s repertoire, and a certain number of demonstrations were needed to secure the association between action and reward.

The establishment of the red box-pushing behavioural variant as a tradition in both experimental single-demonstrator experiments *and* in the free-form multiple-demonstrator experiment further serves to confirm the involvement of social learning. When bees opened boxes in control diffusions, their preference was for the blue behavioural variant (albeit not as pronounced as in colonies exposed to a blue-trained demonstrator). This is not surprising: red sits at the very edge of the bee visual spectrum, while blue is among their most preferred colours (28,29). This strength of preference for blue over red suggests that that naïve bees should be drawn disproportionately towards the blue tab, simply because its colour is a highly salient feature to them. However, in the single-demonstrator diffusions, bees exposed to a demonstrator trained to perform the red-pushing behavioural variant were no less proficient than those exposed to a one that performed the alternative and their preference for their demonstrators’ behavioural variant was no weaker. Even when the alternative blue-pushing behavioural variant was discovered by these learners, they still reverted to their demonstrator’s red-pushing technique.

In the multiple-demonstrator diffusion experiments, the fact that that the *red*-pushing behavioural variant went to fixation in one of the two populations, even though demonstrations of the blue-pushing behavioural variant were readily available to them, further solidifies this conclusion. Here, most proficient learners acquired a preference for one of the two possible behavioural variants. Once an individual had acquired a preference this seemed to be fixed, and the predominance of red- and blue-pushing behaviour was largely governed by stochasticity associated with experienced individuals retiring from foraging and new learners arising. As with all stochastic processes, it is possible that in larger groups the establishment of stable local traditions would be slower or even attenuated. However, these dynamics mirror those of seen in humans, where cultural shifts sometimes come about by individuals with long-held views passing away, rather than changing their personal opinions (34).

While the exact mechanisms underlying the development of individual behavioural preferences in bumblebees remain unclear and warrant further investigation, it does seem likely that subsequent learners in this experiment would have continued to acquire a preference for the red-pushing behavioural variant. The strength of bias towards this variant on days 10-12, in experiment 3, was comparable to that observed in the experimental single-demonstrator diffusions, and learners in these experiments robustly acquired their demonstrator’s preference. Thus, this could be considered to represent a stable local tradition, at least in theory (and under laboratory conditions).

Whether bumblebees are likely to develop similar behavioural traditions naturally remains an open question. The lifespan of *Bombus terrestris* workers is typically only a few weeks from the point of emergence from the pupae, and colonies collapse once the new queens have departed (35,36). Although there may be multiple, sequential sets of workers present during that time, these do not represent true biological generations, and if no workers survive past the decline of annual colonies at the end of the season, any foraging traditions should be lost with them. Thus, it seems unlikely that *Bombus terrestris* would build cultures that span biological generations in the wild, but the results reported here support the notion that the cognitive capacities for this to occur are certainly in place. The Baldwin effect might provide a more roundabout route for culture spanning biological generations in bumblebees, if selection favours those queens whose worker progeny are more likely to develop such traditions. However, there are species of social insect that form colonies that last for years. These include honeybees (37), certain tropical bumblebee species including *Bombus medius* (38), *Bombus atratus* (39), *Bombus rufipes* (40), and stingless bees (41,42). If the learning abilities of these species resemble those of *Bombus terrestris*, it seems plausible that such culture might be found naturally among them.

Even if culture is rare or non-existent in extant wild social insects, the mere existence of this capacity in an invertebrate (given the right conditions and opportunity, as provided here) makes it plausible that cultural processes have in the past contributed to the emergence of behavioural innovations regarding foraging specialisations, nesting architecture and colony organisation. Perhaps elements of the vast, complex repertoire of innate behaviours seen in social insects were not always so instinctive (21). The reason that we have often failed to see evidence of culture in non-human animals may be that we are simply looking too late (5). Social insects may represent an exciting model to investigate these hypotheses, as well as the mechanisms underpinning culture itself.

## Materials and methods

### Animal model

Colonies of bumblebees (*Bombus terrestris audax*) were obtained from Agralan, Ltd. (Swindon, UK) or Koppert Biological Systems Nederland (Berkel en Rodenrijs, Netherlands). Bees were housed in 30.0 × 14.0 × 16.0 cm bipartite wooden nest boxes, and all individuals were marked with numbered Opalith tags for individual identification during transfer to these nest boxes. This involved trapping each bee in a small cage, gently pressing it against the mesh with a sponge, and affixing the tag to the dorsal thorax with a small amount of glue. The nest boxes were connected to flight arenas (66.0 × 60.0 × 30.0 cm or 132.0 × 60.0 × 30.0 cm for single- and multiple-demonstrator experiments, respectively) via 26.0 × 3.5 × 3.5 cm clear acrylic tunnels, which could be blocked to limit access to the flight arena. Bees were allowed to forage *ad libitum* on 20% w/w sucrose solution provided in mass feeders in these arenas overnight, and pollen was provided every two days. Colonies were maintained at standardised room temperature throughout the study, and experiments were conducted under standardised artificial lights (12:12, high-frequency fluorescent lighting; TMS 24F lamps with HF-B 236 TLD [4.3 kHz] ballasts [Koninklijke Phillips NV, Amsterdam, Netherlands]; fitted with Activa daylight fluorescent tubes [OSRAM Licht AG, Munich, Germany]).

### Experimental set-up and puzzle box design

*Puzzle box*. The two-option puzzle box (Fig. 1A) incorporated a rotating, transparent lid that could turn either clockwise or anticlockwise around a central axis, effectively providing two possible ways to “open” the box and obtain a reward; in this case, 50% w/w sucrose solution placed on a yellow “target”. This target was always visible due to the transparent lid, but inaccessible without rotating the lid either clockwise or anti-clockwise by pushing a red tab or a blue tab, respectively. Other than the direction of rotation and tab colour, there was no difference between the two box-opening behavioural variants, which will henceforth be referred to as the “red-pushing behavioural variant” and the “blue-pushing behavioural variant”. A stopper attached to the edge of the boxes prevented bees pushing further onwards after accessing the reward, and a plastic strip around the circumference acted as a “shield” to prevent bees obtaining the reward by reaching with their proboscis from the side.

#### Flight arena

Experiments were conducted in specially-designed flight arenas (see Fig. 1A and Fig. 5). Flaps were cut into the side of the arena through which the puzzle boxes could be removed and replaced, with minimal disturbance to the bees inside the arena. Brush strips lined the inside of the flight arena to prevent bees from escaping during this process. The top of the flight arena was a sheet of transparent UV-transmitting acrylic sheet, and cameras were placed on top of the arena so that the bee ID tags could be captured while recording the diffusion experiments. The flight arenas for the single-demonstrator diffusion experiments had space for 8 puzzle boxes (Fig. 1A), while those used for the multiple-demonstrator diffusion experiments were expanded to provide space for 16 puzzle boxes (Fig. 5). This was done to reduce competition for boxes. To ensure an adequate supply of naïve foragers, the arena was also made bipartite to allow the inclusion of two colonies in a single experimental population. The two nest boxes were connected to opposite ends of the flight arena with a central divider positioned between them. This was removed during the diffusion experiments, but was reinstated to allow the two colonies to forage in separate flight arenas overnight. No inter-colony aggressive interactions were observed during the experiment, and care was taken to return bees to their original colonies after each diffusion session.

#### Demonstrator training protocol

Potential demonstrators were identified during initial group foraging on yellow acrylic chips, which were placed in the flight arena and loaded with 50% w/w sucrose solution. When a bee was observed repeatedly and reliably coming back and forth between the nest box and the flight arena to forage, it was selected for further training. All other bees were restricted to the nest box, and the chosen individual was subjected to an incremental training protocol to learn how to open the puzzle box in one of the two possible ways; either by pushing the red tab clockwise (media file 1) or the blue tab anticlockwise (media file 2; Fig. 1B). The reward used throughout training was 10 μl 50% w/w sucrose solution, and boxes were wiped with 70% ethanol each time they were refilled to remove any olfactory cues.

Training began with the puzzle boxes presented in the “fully open” position, with the yellow target completely exposed and the tab of the selected colour positioned closest to it (Fig. 1Bi shows the starting position for training a bee to push the blue tab). The position of the tab was then modified over time; rotating further and further over the yellow target until it was inaccessible without the bee pushing against the tab. At this point, the bees would inadvertently move the tab forwards while probing the gap beneath it with the proboscis, apparently on an incidental basis. However, as training continued, this behaviour became noticeably more directed. To prevent bees losing motivation, if they failed to access the reward at any point, the reward in the next box was made easier to access.

Training continued until the two tabs were almost equidistant from the yellow target, with the trained tab being ∼1.0 cm closer. At this point the bee progressed to a learning test, which consisted of a box with the two doors equidistant from the yellow target and distilled water in place of the sucrose solution reward. Bees were permitted to leave the flight arena after 5 min, but if the box remained unopened this was considered a failed test. If the box remained unopened after 10 min, the test also ended in failure, and individuals who failed the test were returned to training until they met test criteria again. However, if the bee opened the box successfully within the time limit, it was used as a demonstrator for the open diffusion experiments.

Demonstrators were trained using the same protocol for both the single- and multiple-demonstrator diffusion experiments. The sole difference was that for the multiple-demonstrator diffusion experiments, four demonstrators were trained over two days (two to perform the red-pushing behavioural variant, and two to perform the blue-pushing behavioural variant). The two “red demonstrators” were trained together, and the two “blue demonstrators” were trained together to save time. The reason two demonstrators were trained for each variant was simple: trained demonstrators might, on occasion, die before a diffusion experiment had commenced. As the main point of the experiment was to have both behavioural variants being demonstrated simultaneously with as close to equal incidence as possible at the start, it was decided that two should be trained to perform each variant.

#### Single-demonstrator open diffusion protocol

Colonies seeded with a demonstrator trained to perform the red-pushing behavioural variant will henceforth be referred to as “red colonies”, and those seeded with a demonstrator trained to perform the blue-pushing behavioural variant will be termed “blue colonies”. Experiments were conducted for either six or twelve consecutive days; six days to provide proof of concept, and twelve to see whether box-opening behaviour would persist in a group for longer. Each day at ∼9.30 am, the mass feeders were removed from the flight arena and bees were returned to the nest box. If more than two honeypots were full, the sucrose solution was removed to ensure a strong motivation to forage. After ∼30 min, bees were allowed unrestricted access to the flight arena, where they received 30 min group pre-training with eight lidless boxes, with the yellow targets (bearing 10 μl 50% sucrose solution rewards) fully exposed. The absence of the lid prevented bees from making associations between either tab colour and the reward during this time, which was primarily to ensure that the bees maintained a strong association between the colour yellow and the reward, and to encourage as many into the flight arena as possible before the proper diffusion began (taking advantage of the natural honey-pot monitoring and food alert behaviours of bumblebees (Dornhaus and Chittka, 2001, 2005)). Following this, the boxes were removed, wiped with 70% ethanol, and the targets were refilled with 20 μl 50% w/w sucrose solution. This doubling of the reward volume ensured that the demonstrator would still be sufficiently rewarded if observers began “scrounging”; taking the reward from boxes opened by the demonstrator without opening any themselves. This behaviour was commonplace during pilot studies, so this measure was taken to prevent the demonstrator losing motivation to forage.

The diffusion experiment commenced immediately following pre-training, and the bees were presented with eight fully closed boxes for 3 h. The experimental colonies were seeded with a trained demonstrator while control colonies lacked one, but all other aspects of the protocol were identical. When boxes were depleted of sucrose solution, they were removed from the flight arena through the flaps, washed with 70% ethanol, refilled and then replaced. In the single-demonstrator diffusion experiments, if the demonstrator attempted to perform the behavioural variant it had not been trained on (e.g. attempting to open the box by pushing the blue tab anticlockwise when it was initially trained to push the red tab clockwise) it was prevented from doing so, with the experimenter holding the lid of the box closed with tweezers. No other bees were inhibited from performing either behavioural variant in any way. This was done to try and maintain as much consistency between demonstrators as possible, so the experimental replicates would be more comparable. However, demonstrators attempted the alternate behaviour very rarely (see Appendix Table IA).

A total of six colonies were used for the 6-day diffusions, two of which were seeded with a demonstrator trained to perform the red-pushing behavioural variant (R1, R2), two seeded with a demonstrator trained to perform the blue-pushing behavioural variant (B1, B2) and two control colonies (C1, C2). A further four colonies were used for the 12-day diffusions (B3, R3, C3 and C4), following these naming conventions. In the event that a demonstrator died before any observers met learning criteria, a new bee was selected for individual training. Pre-training and 3 h open diffusion immediately followed a successful learning test. This situation occurred on both day 1 and 2 for colony R3. However, if a demonstrator died after an observer met learning criteria, it was not replaced. The 3 h open diffusions were filmed from above using iPhones (30 fps, 720p; one camera per two boxes to ensure number tags were clear).

#### Multiple-demonstrator open diffusion protocol

The multiple-demonstrator diffusion protocol was identical to the single-demonstrator diffusion protocol, with two exceptions. First, no bees were restricted from performing either behaviour. This would have been too difficult to achieve with four individuals at once, and it was possible that the demonstrators might change their own trained preferences during the experiment as part of a local tradition becoming established (or not). In any case, as in the single-demonstrator diffusion experiments, the demonstrators rarely attempted to perform the alternative, non-trained technique (Appendix Table IB). The second change was that, due to the free-form nature of this experiment and the presence of multiple demonstrators for each variant, demonstrators were not replaced if they died. Two colonies were used for this experiment, and were combined to form a single population as aforementioned.

### Video analysis

Video analysis was conducted in BORIS 7.10.2 (45), and point events were coded whenever a box was opened. In some cases, the tabs would initially be pushed by a bee that would leave before opening the box fully; thus, each opening was assigned to the ID of the bee responsible only when they pushed a tab ≥50% of the required distance to obtain the reward. In cases where observers obtained the reward by pushing <50% of the required distance, these opening instances were included at the overall colony level but were not assigned to an individual; thus, these were absent from observer-specific data. This ensured that all openings assigned to observers involved directed, sustained pushing at the tab, and so unlikely to be by chance.

### Learning criteria

Untrained bees were considered to have made the transition to learners when they had performed full box-opening twice, irrespective of behavioural variant, as this repetition of the behaviour suggested it was not done at random. These criteria split the untrained bees into two groups: *learners* and *non-learners*: those who had met criteria and those who had not.

### Learner proficiency

For the single-demonstrator diffusion experiments, in order to compare learners across colonies, individual proficiency indices (*p*_*i*_) were calculated for each learner as follows:

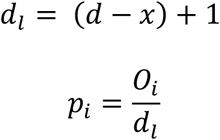

where *O*_*i*_ represented the total opening incidence by the individual in question, *d*_*l*_ represented the total of days spent as a learner (inclusive of the day learning criteria was met), *d* the length of the diffusion in days, and *x* the day learning criteria was met. This allowed comparison between learners that met criteria at different points in the diffusion, e.g. a bee that learned on the second day may have accumulated more openings than a bee that learned on the fifth day not due to proficiency; simply by means of having more time.

As two distinct clusters of learners emerged during the single-demonstrator diffusion experiments: those that performed the target behaviour repeatedly and often, and those that did so rarely and sporadically (in some cases, only performing the behaviour the two times required to meet learning criteria), a simpler method was used to classify learners based on their proficiency in the multiple-demonstrator diffusion experiments. Bees that performed box-opening behaviour (of either variant) >10 times were considered *proficient learners*, while those that performed the behaviour ≤10 times were considered *non-proficient learners*.

### Learner preference

In the multiple-demonstrator diffusion experiments, learners were classed as having a preference for the red variant, a preference for the blue variant, or no preference, based on the proportion of each behavioural variant that they performed throughout the experiment. The boundaries were as follows: red preference, >60% red variant; blue preference, >60% blue variant; no preference, no variant >60% (so, both variants being ≤60% and ≥40%).

### Statistical analysis

Data were analysed with R 4.0.3 (46) and figures were produced using ggplot2 (47). Due to non-normally distributed data, Wilcoxon signed-rank tests were used to compare paired samples and Mann Whitney-U tests were used to compare unpaired samples.

When comparing individual learner proficiency (*p*_*i*_) between “experimental” and “control” colonies, there was no significant difference in *p*_*i*_ between “red colonies” and “blue colonies”, allowing these data to be pooled into a single “experimental” group (W=84.5, p=0.268; Mann Whitney U test). Spearman’s rank order correlation tests were used to analyse the relationship between experimental day and incidence of box-opening by observers.

Two linear mixed-effects models were also used to analyse the data, using the R packages *lme4* (48) and *lmertest* (49). The first analysed the effect of demonstrator presence on learner proficiency over time and included two categorical fixed effects: one between-subjects factor “treatment” (experimental, control) and one within-subjects factor “day” (day 1, day 3; where day 1 was the day an individual met the learning criteria and day 3 was two days following this). Individuals that learned too late in the diffusion to have any data recorded for day 3 were excluded (e.g. a bee that met criteria on day 5 or 6 in the 6-day diffusion experiments, or day 11 or 12 of the 12-day diffusion experiments; n=4 from the experimental colonies and n=3 from the control colonies, leaving n=18 and n=11 in each group, respectively). The response variable was box opening incidence. There was no significant difference between the learners from experimental colonies seeded with “red-pushing demonstrators” and those seeded with “blue-pushing demonstrators” on either day 1 (independent samples Mann Whitney U test; W=55, p=0.1440) or day 3 (independent samples Mann Whitney U test; W=35.5, p=0.4175), so these were pooled into a single experimental group.

The second model analysed the effect of time on individual learner preferences, and included one categorical fixed effect: one within-subjects factor “day’’ (day 1, day *x*; where day 1 was the day an individual met learning criteria and day *x* was the last day they were recorded performing box-opening). Individuals only active on one day (n=1) were excluded, leaving n=12 individuals (n=8 from population 1R2B2 and n=4 from population 2R2B2). The response variable was the proportion of box-opening behaviour that was of the blue-pushing behavioural variant.

We used the Akaike information criterion (AIC) to compare three variations of each model: using bee ID as the sole random factor, using colony/population ID as the sole random factor, and using bee ID nested within colony/population ID, to find the version with the best fit (see Appendix Table 2 and 8). In each case, the model with the lowest AIC was chosen. For all comparisons, P<0.05 was considered to indicate a statistically significant difference.

## Supporting information

Blue Variant

Red Variant

## Acknowledgements

This study was funded by an EPSRC programme grant, “Brains on Board” (ref. no. EP/P006094/1) and a QMUL PhD studentship.

## Author contributions

**A.D.B:** Conceptualization, Methodology, Investigation, Formal Analysis, Writing – Original Draft, Visualisation. **H.M:** Methodology. **O.P:** Methodology. **C.L:** Investigation. **Y.M:** Investigation. **A.K:** Formal Analysis. **J.L.W:** Formal analysis, Writing – Review & Editing. **L.C:** Conceptualization, Funding Acquisition, Writing – Review & Editing.

## Appendix

**Appendix Figure 1.**
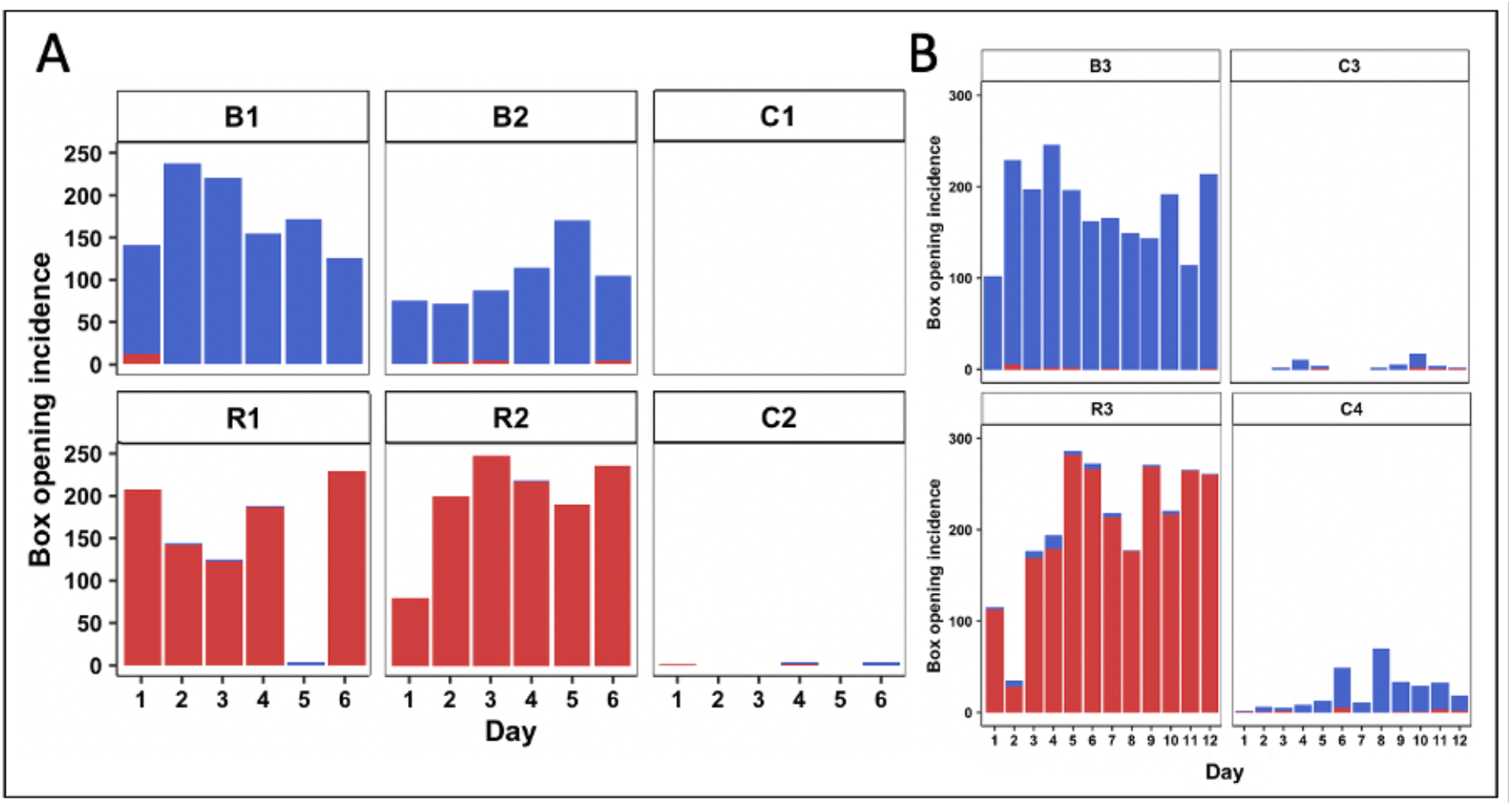
Daily overall box opening incidence in the (A) 6-day and (B) 12-day diffusion experiments. Overall data includes incidences of box-opening by the demonstrator and incidences of box-opening that were not assigned to any observer ID. Colonies B1-3 were each seeded with a demonstrator trained in the blue tab/anticlockwise pushing technique. Colonies R1-3 were each seeded with a demonstrator trained in the red tab/clockwise pushing technique. Colonies C1-4 were controls that lacked a demonstrator. Incidences of the blue tab/anticlockwise pushing technique are depicted in blue, while incidences of the red tab/clockwise pushing technique are depicted in red.

**Appendix Figure 2.**
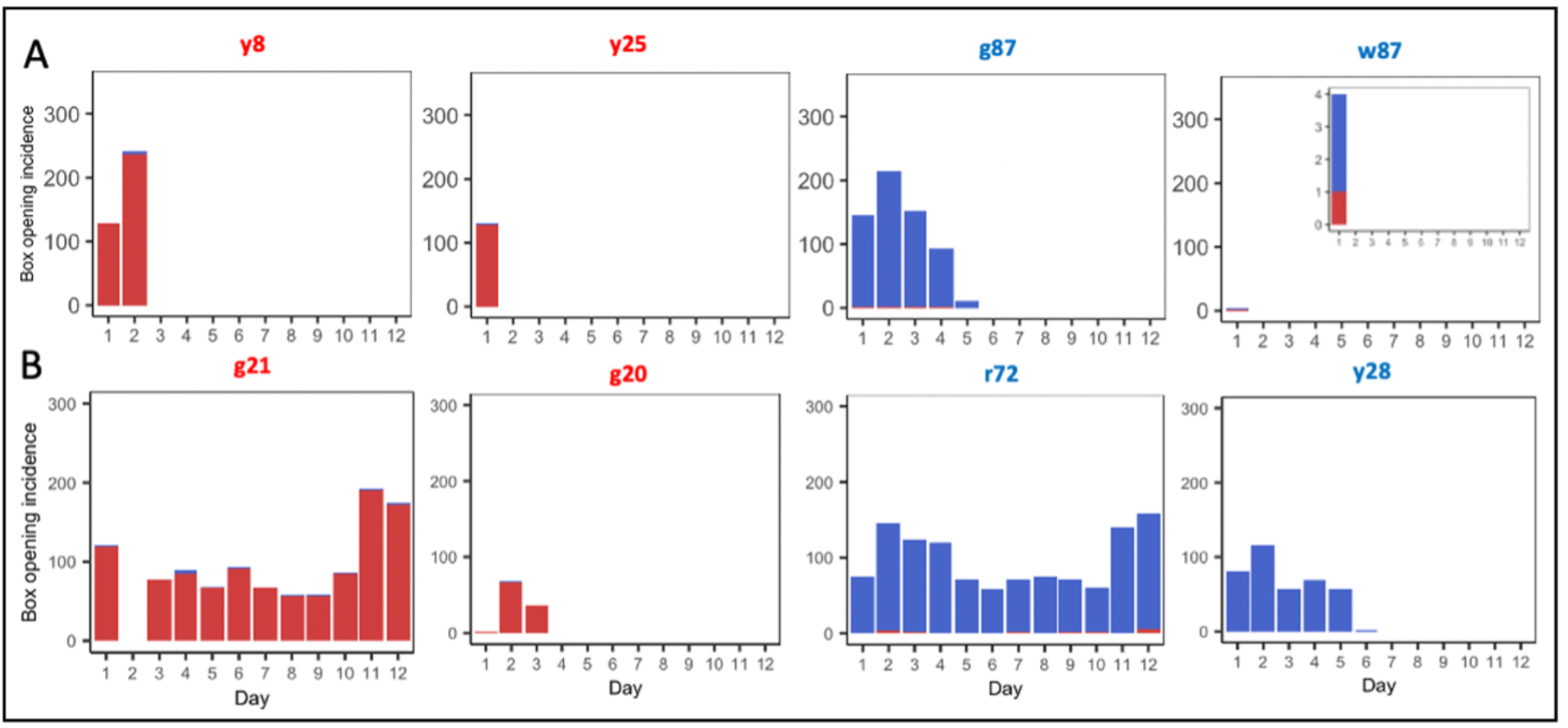
Individual box opening incidence data for demonstrators in (A) population 1R2B2 and (B) population 2R2B2. The colour of the graph titles indicates the trained behavioural variant for each demonstrator. The inset data for w87 is the same as in the wider graph, presented at a finer scale for the sake of clarity.

**Appendix Figure 3.**
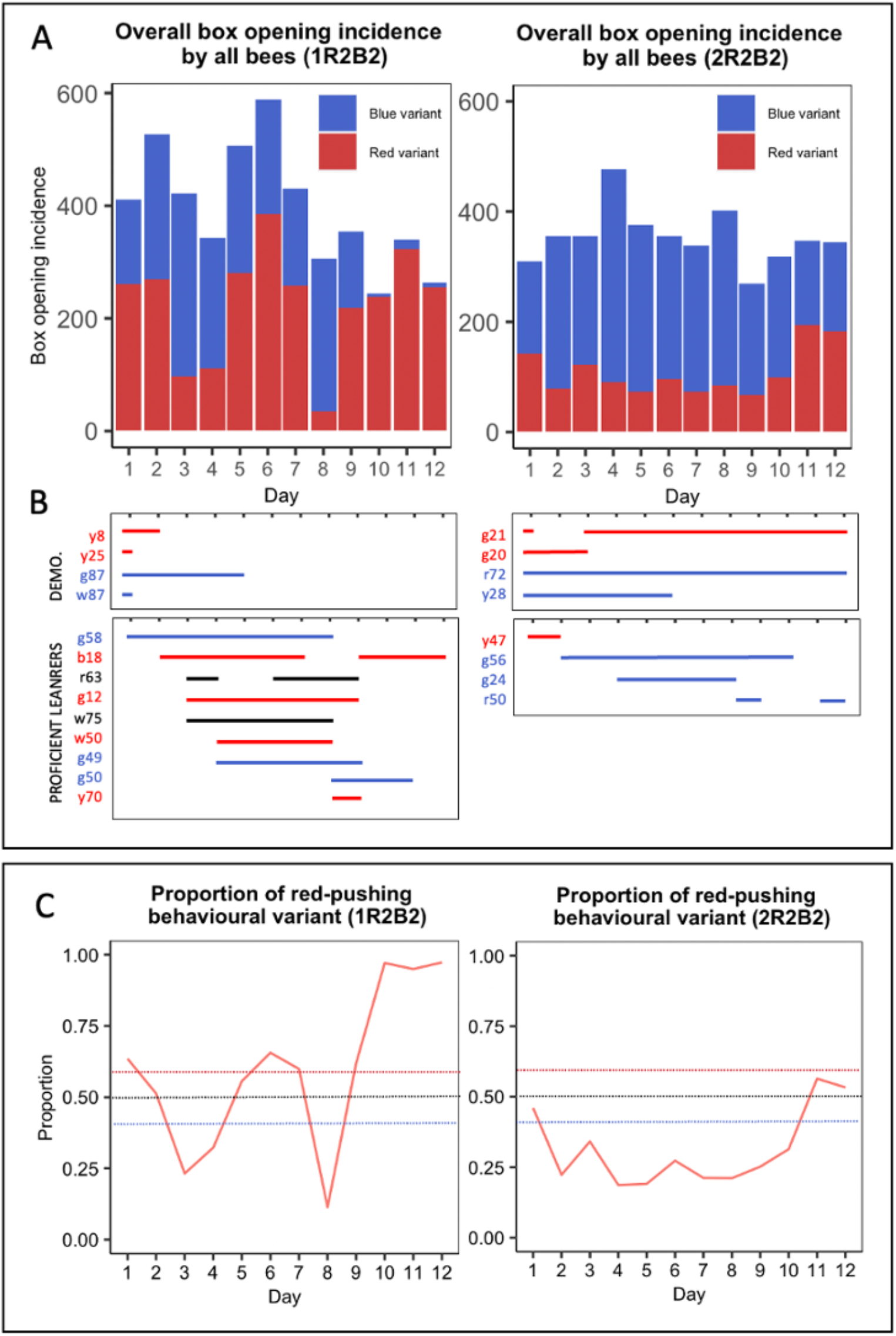
Whole colony data for Experiment 3. Data includes incidences of box-opening by the demonstrator and incidences of box-opening that were not assigned to any observer bee ID. **(A)** Overall box opening incidence by all bees in Experiment 3 (left panel, population 1R2B2, right panel, population 2R2B2). The incidence of each behavioural variant is indicated by colour. **(B)** Days spent active by trained demonstrators and proficient learners. (C) The proportion of recorded daily behaviours that were the red-pushing variant. Dashed lines show the thresholds for a preference for either variant.

**Appendix Table 1.** Total daily box opening and variant incidence (single-demonstrator diffusion experiments)

| A. Overall data<sup>1</sup> |  |  |  |  |  |  |  |  |  |  |  |  |  |  |  |  |  |
| --- | --- | --- | --- | --- | --- | --- | --- | --- | --- | --- | --- | --- | --- | --- | --- | --- | --- |
| Colony ID | Box opening incidence |  |  |  |  |  |  |  |  |  |  |  |  | Total red variant | Total blue variant | Daily average |  |
|  | Day 1 | 2 | 3 | 4 | 5 | 6 | 7 | 8 | 9 | 10 | 11 | 12 | Total |  |  |  |  |
| R1 | 208 | 145 | 124 | 190 | 299 | 229 | - | - | - | - | - | - | 1195 | 1185 | 10 | 199.17 |  |
| R2 | 80 | 199 | 247 | 219 | 190 | 236 | - | - | - | - | - | - | 1171 | 1170 | 1 | 195.17 |  |
| R3 | 115 | 35 | 177 | 194 | 287 | 272 | 218 | 177 | 271 | 220 | 266 | 261 | 2493 | 2438 | 55 | 207.75 |  |
| B1 | 142 | 237 | 220 | 154 | 173 | 126 | - | - | - | - | - | - | 1052 | 14 | 1038 | 175.33 |  |
| B2 | 75 | 72 | 87 | 115 | 170 | 104 | - | - | - | - | - | - | 623 | 12 | 611 | 103.83 |  |
| B3 | 102 | 229 | 197 | 246 | 196 | 162 | 165 | 149 | 144 | 192 | 114 | 214 | 2110 | 12 | 2098 | 175.83 |  |
| C1 | 0 | 0 | 0 | 0 | 0 | 0 | - | - | - | - | - | - | 0 | 0 | 0 | 0.00 |  |
| C2 | 1 | 0 | 0 | 3 | 0 | 3 | - | - | - | - | - | - | 7 | 3 | 4 | 1.17 |  |
| C3 | 0 | 0 | 2 | 11 | 4 | 0 | 0 | 2 | 5 | 17 | 4 | 2 | 47 | 5 | 42 | 3.92 |  |
| C4 | 2 | 6 | 5 | 8 | 13 | 49 | 10 | 70 | 34 | 29 | 33 | 18 | 277 | 16 | 261 | 23.08 |  |
| B. Observer data only<sup>2</sup> |  |  |  |  |  |  |  |  |  |  |  |  |  |  |  |  |  |
| Colony ID | Box opening incidence |  |  |  |  |  |  |  |  |  |  |  |  | Total red variant | Total blue variant | r<sub>s</sub><sup>3</sup> | p |
|  | Day 1 | 2 | 3 | 4 | 5 | 6 | 7 | 8 | 9 | 10 | 11 | 12 | Total |  |  |  |  |
| R1 | 0 | 1 | 35 | 138 | 220 | 171 | - | - | - | - | - | - | 565 | 559 | 6 | 0.94 | 0.017* |
| R2 | 0 | 0 | 0 | 3 | 103 | 113 | - | - | - | - | - | - | 219 | 218 | 1 | 0.94 | 0.005** |
| R3 | 1 | 0 | 4 | 9 | 81 | 124 | 89 | 68 | 174 | 142 | 154 | 160 | 1006 | 990 | 16 | 0.90 | <0.001*** |
| B1 | 28 | 83 | 111 | 52 | 100 | 63 | - | - | - | - | - | - | 437 | 14 | 423 | 0.26 | 0.658 |
| B2 | 0 | 0 | 5 | 19 | 101 | 57 | - | - | - | - | - | - | 182 | 2 | 180 | 0.93 | 0.008** |
| B3 | 0 | 7 | 1 | 3 | 1 | 3 | 164 | 149 | 143 | 191 | 109 | 209 | 980 | 9 | 971 | 0.81 | 0.001** |
| C1 | 0 | 0 | 0 | 0 | 0 | 0 | - | - | - | - | - | - | 0 | 0 | 0 | - | - |
| C2 | 0 | 0 | 0 | 3 | 0 | 2 | - | - | - | - | - | - | 5 | 2 | 3 | 0.54 | 0.268 |
| C3 | 0 | 0 | 2 | 11 | 3 | 0 | 0 | 0 | 4 | 16 | 4 | 1 | 41 | 4 | 37 | 0.30 | 0.347 |
| C4 | 2 | 4 | 5 | 11 | 16 | 43 | 9 | 69 | 34 | 27 | 32 | 17 | 270 | 16 | 254 | 0.70 | 0.015* |

<sup>1</sup>Overall data includes incidences of box-opening by the demonstrator and incidences of box-opening that were not assigned to any untrained bee ID. <sup>2</sup>Box openings were assigned to observers only when they pushed a tab $\geq 50\%$ of the required distance to open the box; and obtained the reward. <sup>3</sup>For the 6-day diffusions, df=4. For the 12-day diffusions, df=10. \*p<0.05, \*\*p<0.01 and \*\*\*p<0.001. Data were analysed using Spearman's rank order correlation tests.

**Appendix Table 2.**
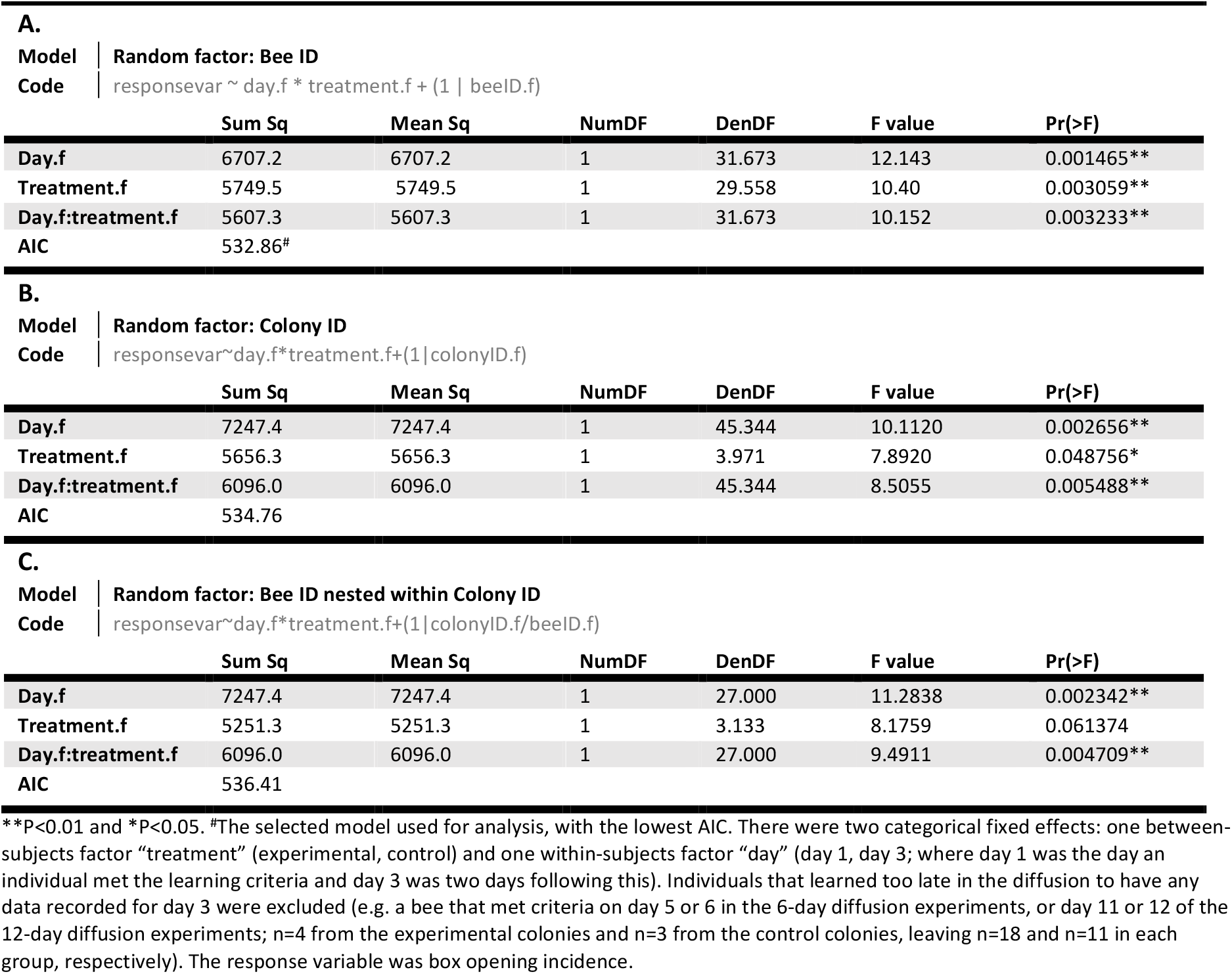
Results of linear mixed-effects model to assess the effect of demonstrator presence on learning proficiency over time.

**A.**
| **Model** | **Random factor: Bee ID** |  |  |  |  |  |
| --- | --- | --- | --- | --- | --- | --- |
| **Code** | responsevar ~ day.f * treatment.f + (1 \| beeID.f) |  |  |  |  |  |
|  | Sum Sq | Mean Sq | NumDF | DenDF | F value | Pr(>F) |
| Day.f | 6707.2 | 6707.2 | 1 | 31.673 | 12.143 | 0.001465** |
| Treatment.f | 5749.5 | 5749.5 | 1 | 29.558 | 10.40 | 0.003059** |
| Day.f:treatment.f | 5607.3 | 5607.3 | 1 | 31.673 | 10.152 | 0.003233** |
| AIC | 532.86<sup>#</sup> |  |  |  |  |  |

**B.**
| **Model** | **Random factor: Colony ID** |  |  |  |  |  |
| --- | --- | --- | --- | --- | --- | --- |
| **Code** | responsevar~day.f*treatment.f+(1\|colonyID.f) |  |  |  |  |  |
|  | Sum Sq | Mean Sq | NumDF | DenDF | F value | Pr(>F) |
| Day.f | 7247.4 | 7247.4 | 1 | 45.344 | 10.1120 | 0.002656** |
| Treatment.f | 5656.3 | 5656.3 | 1 | 3.971 | 7.8920 | 0.048756* |
| Day.f:treatment.f | 6096.0 | 6096.0 | 1 | 45.344 | 8.5055 | 0.005488** |
| AIC | 534.76 |  |  |  |  |  |

**C.**
| **Model** | **Random factor: Bee ID nested within Colony ID** |  |  |  |  |  |
| --- | --- | --- | --- | --- | --- | --- |
| **Code** | responsevar~day.f*treatment.f+(1\|colonyID.f/beeID.f) |  |  |  |  |  |
|  | Sum Sq | Mean Sq | NumDF | DenDF | F value | Pr(>F) |
| Day.f | 7247.4 | 7247.4 | 1 | 27.000 | 11.2838 | 0.002342** |
| Treatment.f | 5251.3 | 5251.3 | 1 | 3.133 | 8.1759 | 0.061374 |
| Day.f:treatment.f | 6096.0 | 6096.0 | 1 | 27.000 | 9.4911 | 0.004709** |
| AIC | 536.41 |  |  |  |  |  |

\*\*P<0.01 and \*P<0.05. <sup>#</sup>The selected model used for analysis, with the lowest AIC. There were two categorical fixed effects: one between-subjects factor “treatment” (experimental, control) and one within-subjects factor “day” (day 1, day 3; where day 1 was the day an individual met the learning criteria and day 3 was two days following this). Individuals that learned too late in the diffusion to have any data recorded for day 3 were excluded (e.g. a bee that met criteria on day 5 or 6 in the 6-day diffusion experiments, or day 11 or 12 of the 12-day diffusion experiments; n=4 from the experimental colonies and n=3 from the control colonies, leaving n=18 and n=11 in each group, respectively). The response variable was box opening incidence.

**Appendix Table 3.**
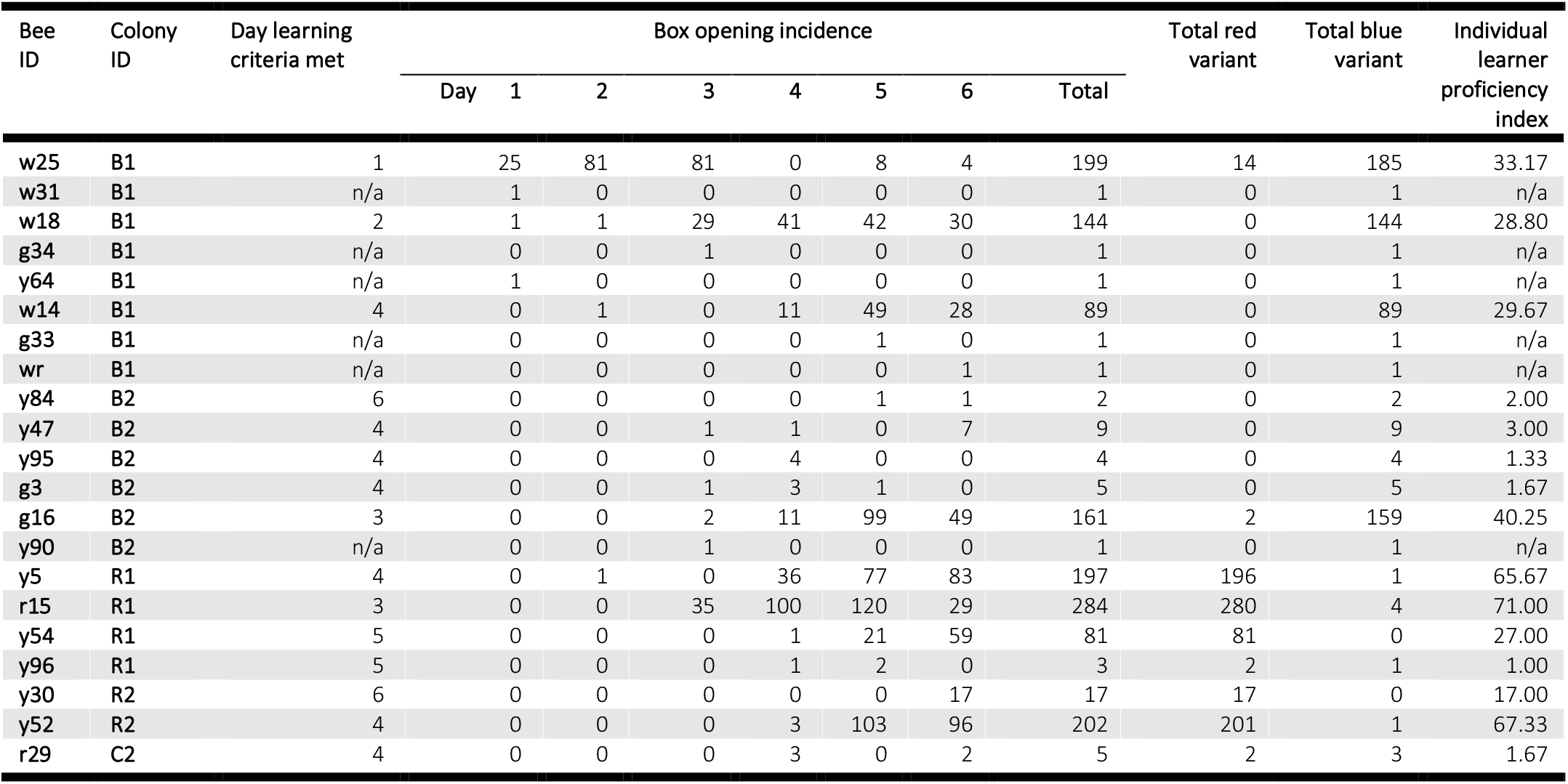
Daily box opening incidence by individual observers (single-demonstrator 6-day diffusion experiments)

**Appendix Table 4.** Daily box opening incidence by individual observers (single-demonstrator 12-day diffusion experiments)

| Bee ID | Colony ID | Day learning criteria met | Box opening incidence |  |  |  |  |  |  |  |  |  |  |  |  | Total red variant | Total blue variant | Indiv. learner prof. index |
| --- | --- | --- | --- | --- | --- | --- | --- | --- | --- | --- | --- | --- | --- | --- | --- | --- | --- | --- |
|  |  |  | Day 1 | 2 | 3 | 4 | 5 | 6 | 7 | 8 | 9 | 10 | 11 | 12 | Total |  |  |  |
| y47 | B3 | 4 | 0 | 0 | 1 | 2 | 0 | 0 | 0 | 0 | 0 | 0 | 0 | 0 | 3 | 2 | 1 | 0.33 |
| y1 | B3 | 2 | 0 | 7 | 0 | 0 | 0 | 0 | 0 | 0 | 0 | 0 | 0 | 0 | 7 | 4 | 3 | 0.63 |
| r14 | B3 | n/a | 0 | 0 | 0 | 1 | 0 | 0 | 0 | 0 | 0 | 0 | 0 | 0 | 1 | 0 | 1 | n/a |
| r24 | B3 | n/a | 0 | 0 | 0 | 0 | 0 | 0 | 0 | 0 | 1 | 0 | 0 | 0 | 1 | 0 | 1 | n/a |
| r35 | B3 | 6 | 0 | 0 | 0 | 0 | 0 | 3 | 164 | 149 | 139 | 120 | 38 | 156 | 769 | 1 | 768 | 109.86 |
| r27 | B3 | n/a | 0 | 0 | 0 | 0 | 1 | 0 | 0 | 0 | 0 | 0 | 0 | 0 | 1 | 1 | 0 | n/a |
| y40 | B3 | 9 | 0 | 0 | 0 | 0 | 0 | 0 | 0 | 0 | 3 | 71 | 71 | 52 | 197 | 0 | 197 | 49.25 |
| b11 | B3 | n/a | 0 | 0 | 0 | 0 | 0 | 0 | 0 | 0 | 0 | 0 | 0 | 1 | 1 | 1 | 0 | n/a |
| w51 | R3 | n/a | 1 | 0 | 0 | 0 | 0 | 0 | 0 | 0 | 0 | 0 | 0 | 0 | 1 | 1 | 0 | n/a |
| w49 | R3 | 3 | 0 | 0 | 4 | 3 | 22 | 30 | 0 | 0 | 0 | 0 | 0 | 0 | 59 | 56 | 3 | 5.90 |
| g93 | R3 | 7 | 0 | 0 | 0 | 0 | 0 | 1 | 2 | 0 | 5 | 8 | 1 | 5 | 22 | 22 | 0 | 3.67 |
| w35 | R3 | 4 | 0 | 0 | 0 | 6 | 59 | 93 | 87 | 59 | 79 | 77 | 69 | 73 | 602 | 591 | 11 | 66.89 |
| g56 | R3 | 8 | 0 | 0 | 0 | 0 | 0 | 0 | 0 | 9 | 90 | 57 | 84 | 82 | 322 | 320 | 2 | 64.40 |
| w57 | C3 | 3 | 0 | 0 | 2 | 11 | 0 | 0 | 0 | 0 | 0 | 0 | 0 | 0 | 13 | 0 | 13 | 1.30 |
| r24 | C3 | 5 | 0 | 0 | 0 | 0 | 3 | 0 | 0 | 0 | 0 | 0 | 0 | 0 | 3 | 0 | 3 | 0.38 |
| b5 | C3 | 9 | 0 | 0 | 0 | 0 | 0 | 0 | 0 | 0 | 2 | 16 | 3 | 1 | 22 | 4 | 18 | 5.50 |
| r42 | C3 | 9 | 0 | 0 | 0 | 0 | 0 | 0 | 0 | 0 | 2 | 0 | 1 | 0 | 3 | 0 | 3 | 0.75 |
| g91 | C4 | 2 | 1 | 1 | 0 | 0 | 0 | 0 | 0 | 0 | 0 | 0 | 0 | 0 | 2 | 1 | 1 | 0.18 |
| g63 | C4 | 10 | 1 | 0 | 0 | 0 | 0 | 0 | 0 | 0 | 0 | 1 | 0 | 0 | 2 | 2 | 0 | 0.67 |
| g26 | C4 | 2 | 0 | 3 | 0 | 0 | 0 | 0 | 0 | 0 | 0 | 0 | 0 | 0 | 3 | 0 | 3 | 0.27 |
| y46 | C4 | 3 | 0 | 0 | 4 | 3 | 6 | 0 | 1 | 0 | 1 | 0 | 0 | 0 | 15 | 5 | 10 | 1.50 |
| y28 | C4 | n/a | 0 | 0 | 1 | 0 | 0 | 0 | 0 | 0 | 0 | 0 | 0 | 0 | 1 | 0 | 1 | n/a |
| y47 | C4 | n/a | 0 | 0 | 0 | 1 | 0 | 0 | 0 | 0 | 0 | 0 | 0 | 0 | 1 | 0 | 1 | n/a |
| y12 | C4 | 4 | 0 | 0 | 0 | 7 | 10 | 39 | 8 | 68 | 31 | 25 | 21 | 7 | 216 | 2 | 214 | 24.00 |
| g40 | C4 | n/a | 0 | 0 | 0 | 0 | 0 | 1 | 0 | 0 | 0 | 0 | 0 | 0 | 1 | 0 | 1 | n/a |
| g41 | C4 | 6 | 0 | 0 | 0 | 0 | 0 | 3 | 0 | 1 | 0 | 1 | 0 | 1 | 6 | 1 | 5 | 0.86 |
| y52 | C4 | n/a | 0 | 0 | 0 | 0 | 0 | 0 | 0 | 0 | 0 | 0 | 0 | 1 | 1 | 0 | 1 | n/a |
| 00 | C4 | n/a | 0 | 0 | 0 | 0 | 0 | 0 | 0 | 0 | 1 | 0 | 0 | 0 | 1 | 0 | 1 | n/a |
| g24 | C4 | 11 | 0 | 0 | 0 | 0 | 0 | 0 | 0 | 0 | 1 | 0 | 3 | 6 | 10 | 2 | 8 | 5.00 |
| y1 | C4 | 11 | 0 | 0 | 0 | 0 | 0 | 0 | 0 | 0 | 0 | 0 | 2 | 0 | 2 | 1 | 1 | 1.00 |
| g18 | C4 | 11 | 0 | 0 | 0 | 0 | 0 | 0 | 0 | 0 | 0 | 0 | 6 | 2 | 8 | 2 | 6 | 4.00 |

**Appendix Table 5.** Demonstrator characteristics in the multiple-demonstrator diffusion experiments.

| Bee ID | Trained variant | Total incidence | Total red variant incidence | Total blue variant incidence | % Red variant | % Blue variant | Activity period (days) |
| --- | --- | --- | --- | --- | --- | --- | --- |
| *Population 1R2B2* |  |  |  |  |  |  |  |
| y8 | R | 369 | 366 | 3 | 99.19 | 0.81 | 1-2 |
| y25 | R | 131 | 129 | 2 | 98.47 | 1.53 | 1 |
| g87 | B | 615 | 7 | 608 | 1.14 | 98.86 | 1-5 |
| w87 | B | 4 | 1 | 3 | 25.00 | 75.00 | 1 |
| *Overall* |  | *1119* | *503* | *616* | *44.95* | *55.05* |  |
| *Population 2R2B2* |  |  |  |  |  |  |  |
| g21 | R | 1079 | 1067 | 12 | 98.9 | 1.1 | 1, 3-12 |
| g20 | R | 106 | 105 | 1 | 99.1 | 0.9 | 1-3 |
| r72 | B | 1165 | 17 | 1148 | 1.5 | 98.5 | 1-12 |
| y28 | B | 380 | 0 | 380 | 0.0 | 100.0 | 1-6 |
| *Overall* |  | *2730* | *1189* | *1541* | *43.6* | *56.4* |  |

**Appendix Table 6.**
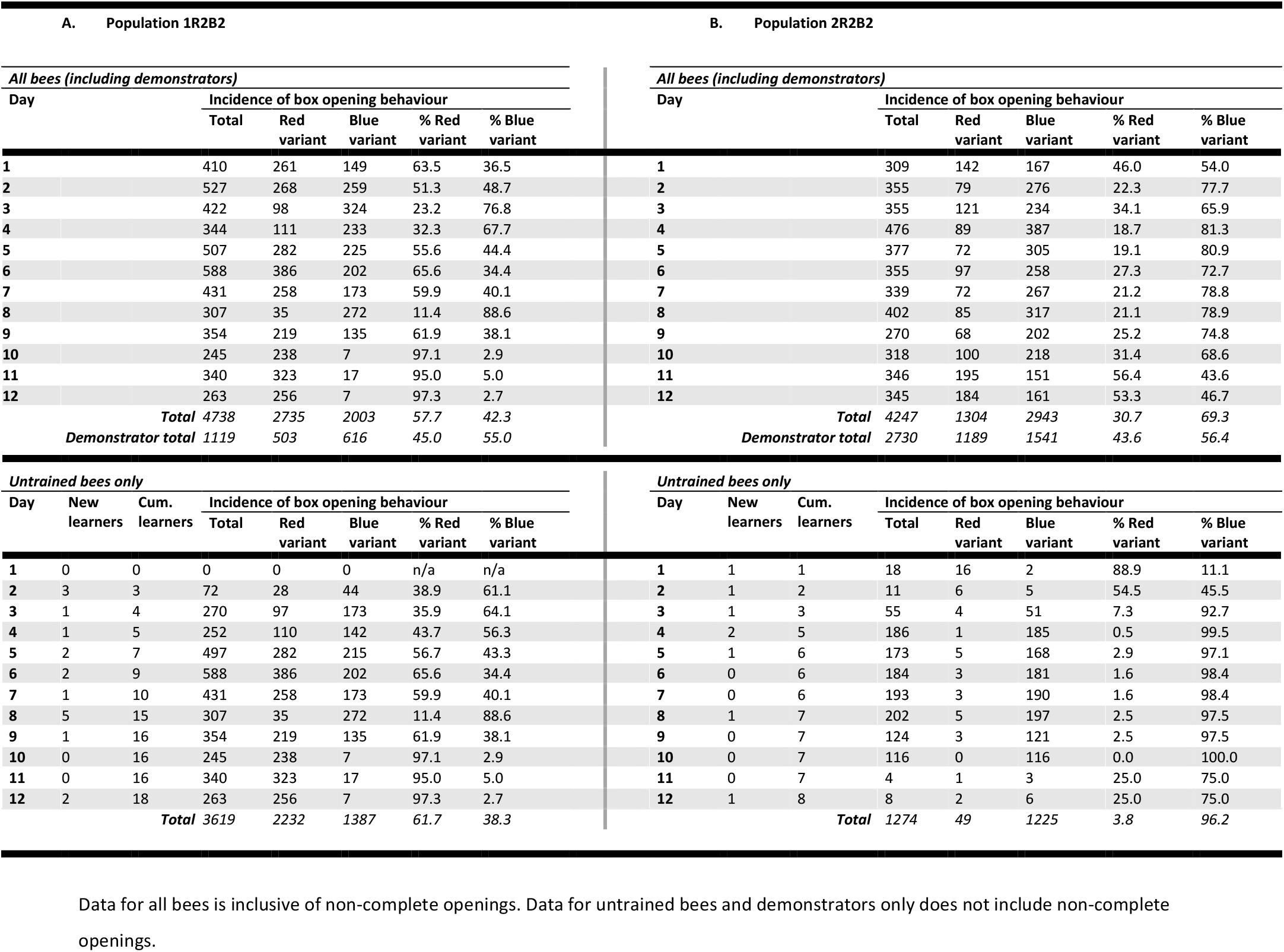
Total daily box opening and variant incidence (multiple-demonstrator diffusion experiments)

**Appendix Table 7.**
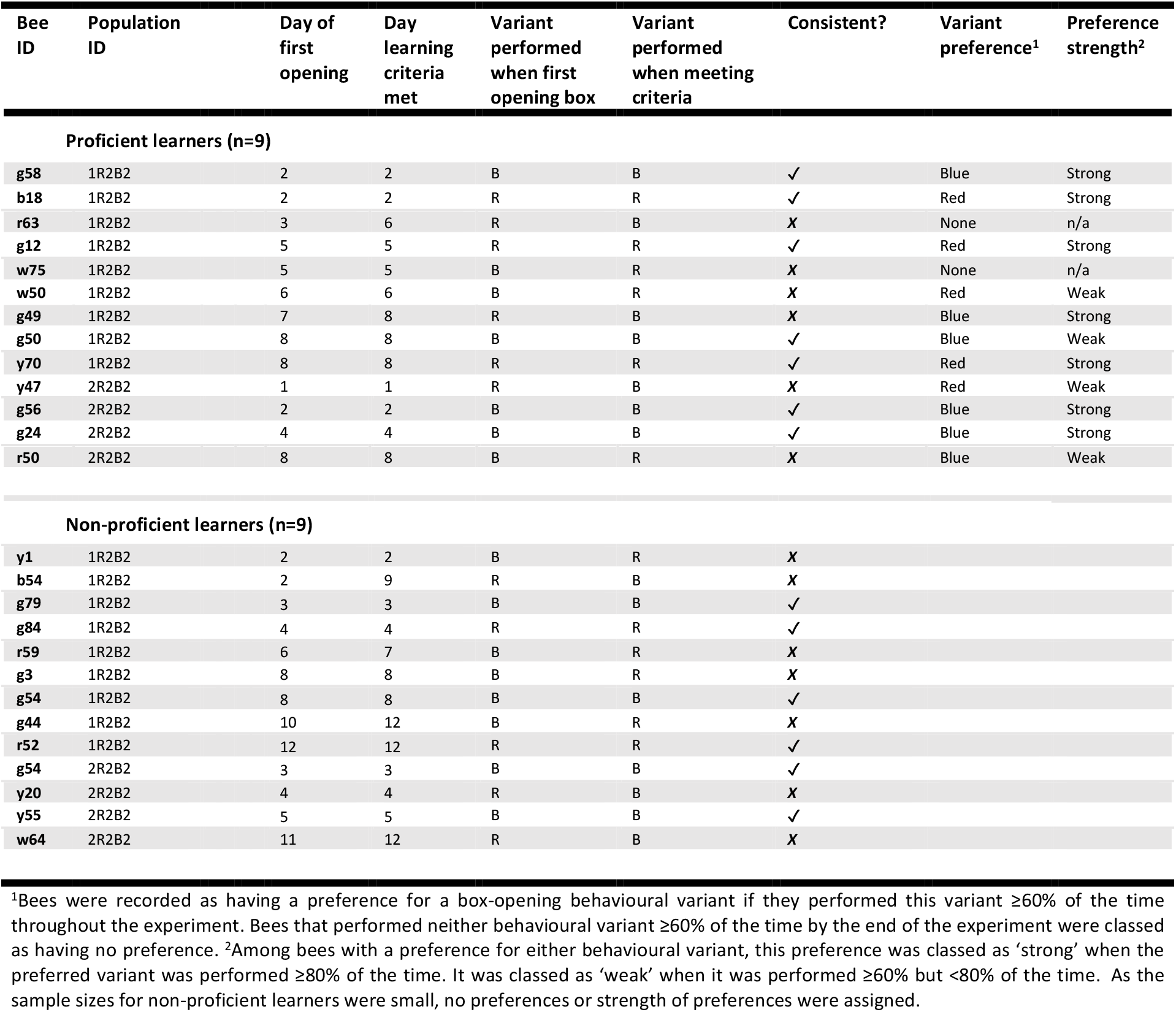
Observer characteristics (multiple-demonstrator diffusion experiments)

**Appendix Table 8.**
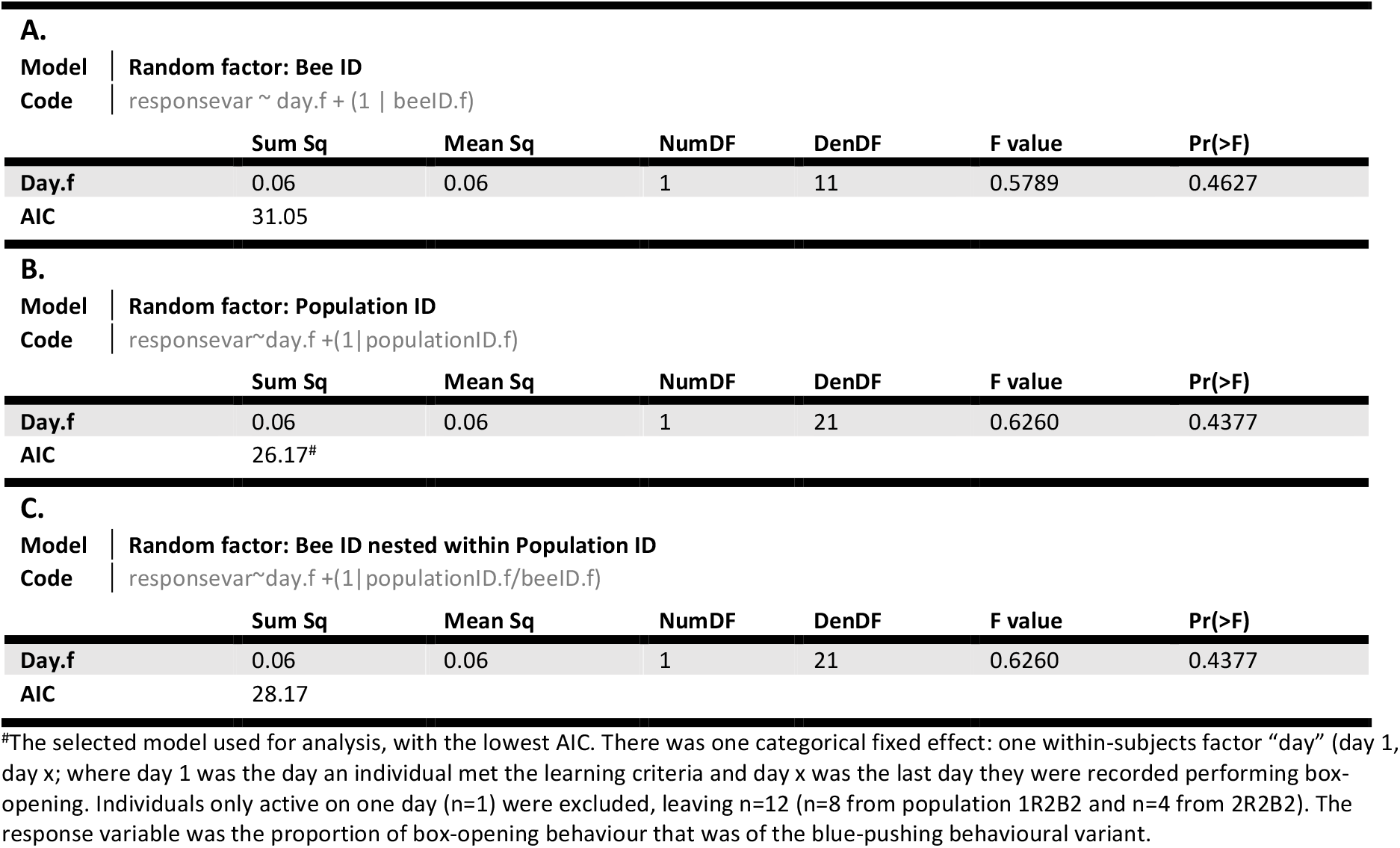
Results of linear mixed-effects model to assess the effect of time on learner preference.

## Supporting information

***Video 1.*** The red-pushing behavioural variant. A bee opens a puzzle box by pushing against the red tab to rotate the lid of the box clockwise (bluevariant.mp4).

***Video 2.*** The blue-pushing behavioural variant. A bee opens a puzzle box by pushing against the blue tab to rotate the lid of the box anticlockwise (redvariant.mp4).

